# Unbiased estimation of the coefficient of determination in linear models: an application to fMRI encoding model comparison

**DOI:** 10.1101/2024.03.04.583270

**Authors:** Agustin Lage Castellanos, Federico De Martino, Giancarlo Valente

## Abstract

Neuroscientific investigation has greatly benefited from the combination of functional Magnetic Resonance Imaging (fMRI) with linearized encoding, which allows to validate and compare computational models of neural activity based on neuroimaging data. In linearized encoding, a multidimensional feature space, usually obtained from a computational model applied to the stimuli, is related to the measured brain activity. This is often done by mapping such space to a dataset (training data, or *in-sample*), and validating the mapping on a separate dataset (test data, or *out-of-sample*), to avoid overfitting. When comparing models, the one with the highest explained variance on the test data, as indicated by the coefficient of determination (*R*^2^), is the one that better reflects the neural computations performed by the brain. An implicit assumption underlying this procedure is that the *out-of-sample R*^2^ is an unbiased estimator of the explanatory power of a computational model in the population of stimuli, and can therefore be safely used to compare models. In this work, we show that this is not the case, as the *out-of-sample R*^2^ has a negative bias, related to the amount of overfitting in the training data. This phenomenon has dramatic implications for model comparison when models of different dimensionalities are compared. To this aim, we develop an analytical framework that allows us to evaluate and correct biases in both *in-* and *out-of-sample R*^2^, with and without L2 regularization. Our proposed approach yields unbiased estimators of the population *R*^2^, thus enabling a valid model comparison. We validate it through illustrative simulations and with an application to a large public fMRI dataset.

## 1 Introduction

Neuroscientific investigation has greatly benefited from validating (neural) computational models using brain responses obtained with neuroimaging techniques, such as functional Magnetic Resonance Imaging (fMRI) [De Martino et al., 2018, Kriegeskorte and Douglas, 2019, Cohen et al., 2017].

In particular, in linearized encoding [Naselaris et al., 2011], a multidimensional feature space describing how stimuli are represented by a computational model, is linearly mapped onto the evoked brain responses. By examining how well different feature spaces (*e*.*g*. different computational models or, in the case of deep neural networks, different layers of the same architecture) explain the data, model comparison can be carried out providing insight into different neural theories [Storrs et al., 2021, Gifford et al., 2023, de Heer et al., 2017].

This approach is now widespread and has resulted in findings across several (perceptual) domains. For example, when considering visual perceptual processes, it has been shown that the hemodynamic responses in the primary visual cortex can be best modeled using low-level image characteristics, based e.g. on a Gabor wavelet decomposition of the input images [Naselaris et al., 2015]. It has additionally been shown that, when moving across the ventral stream, more complex neural computational models better explain the imaging data [Güçlü and van Gerven, 2014, Storrs et al., 2021, Cichy et al., 2016] and that postural features are encoded in regions that have been characterized as selective to body stimuli [Marrazzo et al., 2023]. In the auditory domain, encoding models have been successfully used to characterize the spectral content of natural sounds [Moerel et al., 2012, Moerel et al., 2013], compare models of acoustic processing [Santoro et al., 2014, Norman-Haignere and McDermott, 2018], evaluate models of pitch perception [De Angelis et al., 2018], test deep networks trained for specific tasks [Kell et al., 2018], map the sensitivity to spectral information and sound location in subcortical areas [De Martino et al., 2013] and evaluate the sensitivity of cortical responses to acoustic, articulatory and semantic features of speech [de Heer et al., 2017].

The linearized encoding pipeline consists of several steps. First, the stimuli are described by one or more computational models. In this step, each stimulus is associated with a multidimensional feature vector representing the properties of the stimulus in the space described by the computational model. For this representation to be useful in an encoding analysis, it is important to sample a large set of different stimuli. To this avail, several efforts have been devoted towards the creation of large datasets to be used for testing the encoding of computational models in fMRI [Allen et al., 2022, LeBel et al., 2023]. Next, a link between brain activity (e.g. an fMRI voxel or group of voxels) and a given model is estimated (we will refer to this as the mapping or encoding step to differentiate it from the computational model used to represent the stimuli), and subsequently validated. The estimation part is often carried out by multiple regression, using Ordinary Least Square (OLS) or techniques that better handle high dimensional models (e.g. Ridge Regression) [Hastie et al., 2001]. In the validation step, the estimated mapping is used to predict the brain activity elicited by test stimuli, starting from their computational representation. This is often done using a separate (test) dataset (that includes different stimuli from the ones used in training), similar to what is commonly done in the machine learning literature [Bishop, 2007].

Different metrics have been proposed in the literature to compute the accuracy of the mapping between a computational model and brain responses, such as the correlation coefficient between the observed and predicted brain responses across the different stimuli [Kay et al., 2008, Lage-Castellanos et al., 2019], or the variance explained by a model in the test data (coefficient of determination, *R*^2^) [Gifford et al., 2023, Storrs et al., 2021]. When comparing competing models, the use of *R*^2^ relates to the problem of partitioning the variance explained by each competing model, decomposing it into unique and shared contributions (see commonality analysis [Seibold and McPhee, 1979] and for applications e.g. [de Heer et al., 2017], [Dupré la Tour et al., 2022]). Whereas this decomposition provides a more informed picture of the interplay between the different computational models, determining the best model does not rely on this decomposition as the model with the highest unique contribution is also the one with the highest *R*^2^ value, as we will show in section 3.2.

The coefficient of determination *R*^2^ is commonly interpreted as the proportion of the variance of the brain response that can be explained by the computational model. However, this interpretation is not entirely valid in all cases. As we will show in this article, the coefficient of determination is equal to the ratio between the variance explained by the model and the total variance of the data only when the mapping between brain measures and stimulus representations is estimated using OLS and *R*^2^ is evaluated on the training data.

A well-known property of *R*^2^ in regression problems, such as linearized encoding, is that when it is computed on the same data used to estimate the regression weights (*i*.*e*. “in-sample”), it suffers from an optimistic bias. To avoid this confound, and following common practices used in the machine learning community, it has been advocated, in the fMRI encoding literature, to perform model comparison exclusively on an independent dataset (*i*.*e*. “out-of-sample”). This can either be done with a left-out dataset, or with cross-validation. However, whereas much is known about the in-sample bias (e.g. its dependence on the number of dimensions of the model, and the methods to adjust for it [Ezekiel, 1930a, Olkin and Pratt, 1958, Yin and Fan, 2001]), the bias present in the out-of-sample estimator of the explained variance has received comparatively less attention [Kromrey and Hines, 1995, Chen and Qi, 2023]. Empirical techniques, such as cross-validation, that are commonly used for estimating the out-of-sample *R*^2^, implicitly (and erroneously) assume that the resulting estimate is a valid measure to generalize to the population. In this work, we develop an analytical approach that can be used to evaluate the sources of bias in the in- and out-of-sample *R*^2^. Furthermore, this framework allows the development of unbiased estimators of the population *R*^2^ in both cases and also in the presence of regularization with Ridge Regression. This has vast implications for model comparison by means of explained variance as in *e*.*g*. neuroimaging encoding applications.

To illustrate the consequences of the bias in *R*^2^ we can describe it by means of an example. Consider the case of comparing two computational models (*M*_*A*_ and *M*_*B*_) consisting of *p*_*A*_ = 10 and *p*_*B*_ = 30 (uncorrelated) dimensions and a training set size of 150 samples. In this simulation, the explained variances are, according to the generative model, 21.3% and 28.6% respectively (see Figure 1, and Section 2.7 for more details on the data generation). We further assume that the test data is considerably larger, allowing a precise estimate of the explained variance. In the context of fMRI encoding, this toy example corresponds to comparing the performance (in predicting the fMRI response of a voxel) of two computational models representing 150 stimuli in two feature spaces (*M*_*A*_ and *M*_*B*_) with different number of dimensions. Since the number of features is smaller than the training sample size, we consider using OLS. When fitting each feature space to the data, the variance explained in the training data is 25.6% and 41.3% respectively. This reflects the well-known (optimistic) in-sample bias which is larger for model B due to its higher dimensionality. In this example, performing model comparison based on in-sample *R*^2^ would result in a correct conclusion (i.e. model *M*_*B*_ has higher accuracy than model *M*_*A*_), but with an overestimation of the magnitude of the difference between the variance explained by the two models. Using test data, as commonly done in the encoding literature, does not solve the problem. In our example, the observed *R*^2^ in test data are 14.2% and 9.4% for models *M*_*A*_ and *M*_*B*_ respectively, indicating that a bias is still present, which in this case induces also to erroneously select model (A) as the best model.

**Figure 1:**
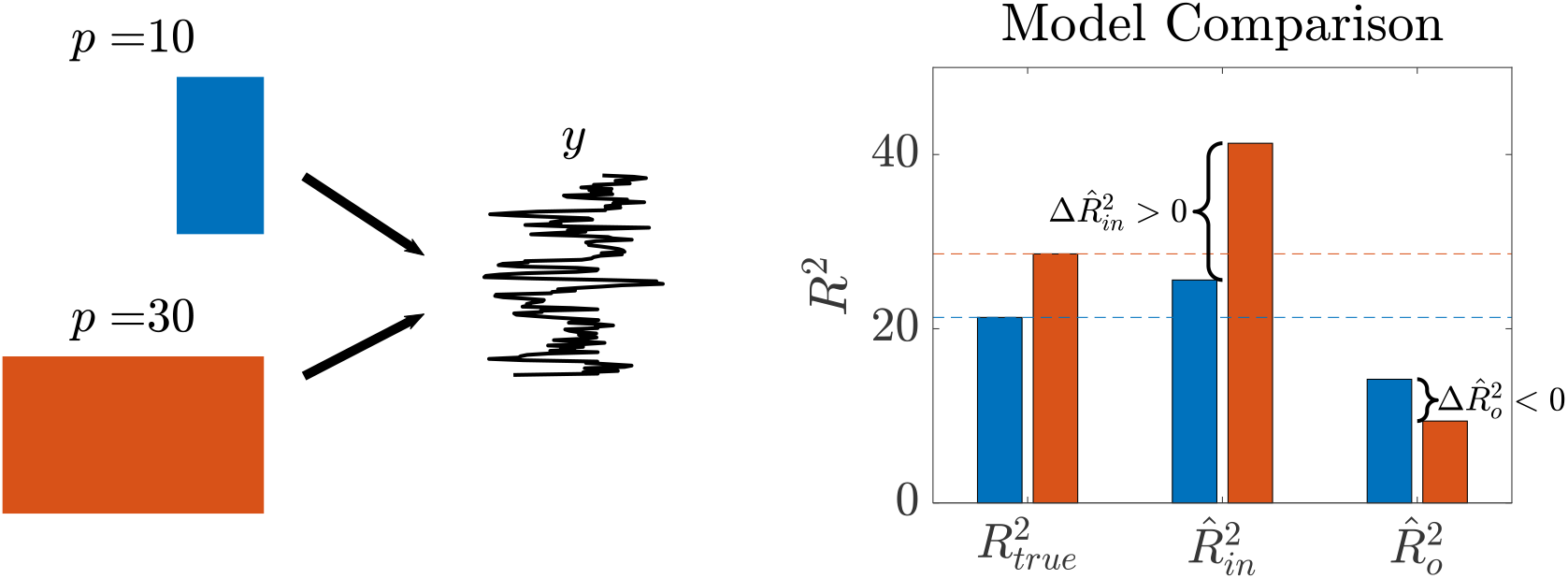
Example of the effect of the bias in *R*^2^ on model comparison. Two feature spaces of different dimensions (*M*_*A*_ with 10 features in blue, and *M*_*B*_ with 30 features in red) are compared on the basis of the variance they explain from the observed response *y* (left panel). The true *R*^2^, the *R*^2^ estimated in the training data 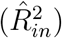 and in the test data 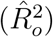 are reported in the right panel. The figure illustrates the possibly contradictory conclusions obtained when comparing models based on in-sample and out-of-sample estimates.

In this work we develop an analytical decomposition of the bias terms that are encountered when estimating *R*^2^ in-sample or out-of-sample, relating both to the phenomenon of overfitting. We furthermore characterize the effect that regularization, with Ridge Regression, has on the estimation of *R*^2^ and we show that two different bias terms arise, that can be linked separately to overfitting and shrinkage. Based on this analytical derivation, we propose a general correction that accounts for the bias terms, resulting in an unbiased estimator of *R*^2^ both in- and out-of-sample. This correction is based on an approximation, and we validate it using simulations and a (large) publicly available fMRI dataset [Allen et al., 2022].

## 2 Methods

### 2.1 Glossary

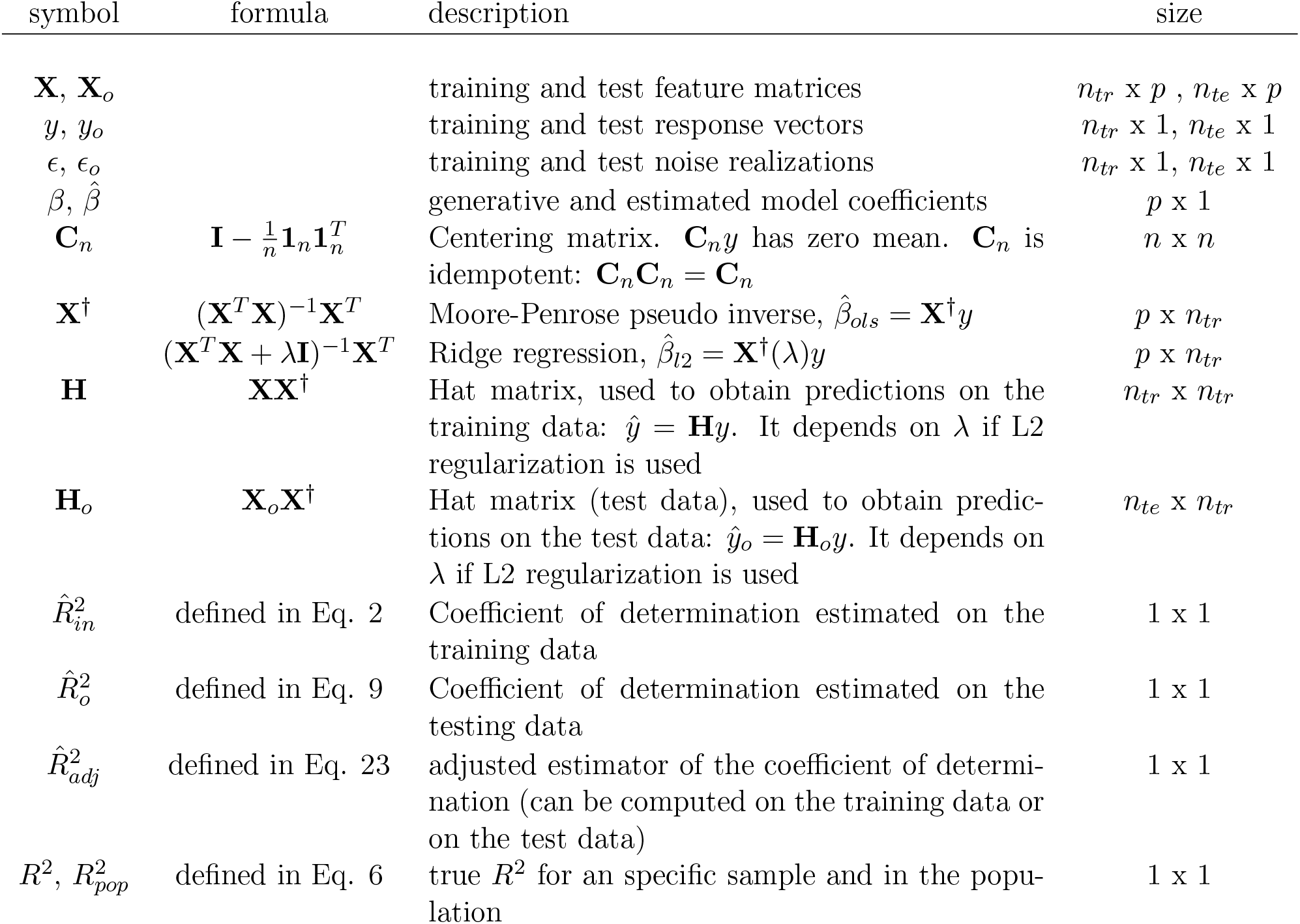

### 2.2 Definition of *R*^2^ and its estimators

Linearized encoding (as applied to e.g. fMRI data) requires the definition of the feature matrix **X** describing how a given set of *n* stimuli is represented by a computational model with *p* features. To understand how the features are encoded in the brain responses, a linear mapping of the feature matrix onto the vector *y* representing the brain responses to the *n* stimuli (measured e.g. with fMRI) is estimated. The goodness of the estimated mapping is often evaluated using the coefficient of determination. We will first define the coefficient of determination, and then characterize its statistical properties based on the generative model that underlies the use of linear regression in this context. The generative model is:

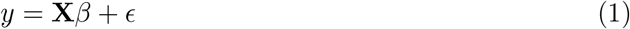

The coefficients *β* represent the mapping between model features and observations and the noise term *ϵ* is assumed to be independent and identically distributed (*i*.*i*.*d*) with variance 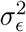. An assumption made when using linear regression is the independence between the noise and the columns of **X** [Rao, 1973]. Additionally, we assume in this work that **X** is a random variable, with a given distribution.

When possible, ordinary least squares (OLS) is used, leading to 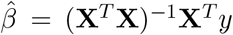. In case of (near) collinearity across the features or when the number of stimuli is smaller than the number of features, regularization can be used to obtain a reliable estimate. Across all variants of regularized least squares, the most frequently used in linearized encoding is ridge regression [Hastie et al., 2001], where 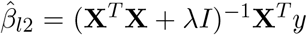, where *λ* is the regularization parameter. The performance of a computational model is usually assessed by computing the *coefficient of determination*, computed as one minus the ratio between the variance of what cannot be explained by the model (Mean Square Error, MSE) and the total variance of the observation *y* (Total Sum of Squares, TSS).

In the linear regression literature, the coefficient of determination was introduced as an *in-sample* measure, where **X** and *y* are the same used to estimate 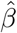. In this case, the fit of the observation is 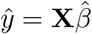and the estimated residuals are 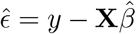. For the sake of conciseness, we avoid providing special emphasis on the model intercept, and thus assume that, if required, it can be represented as a column of constant values within **X**. The estimator of the *R*^2^ in the training data is therefore:

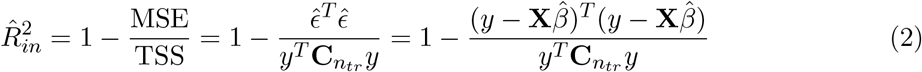

When OLS is used, the two terms 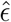 and 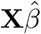 are orthogonal by construction 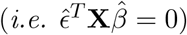 and the mean of the residuals 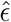 is zero (therefore 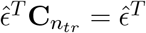). Therefore Eq. (2) simplifies as:

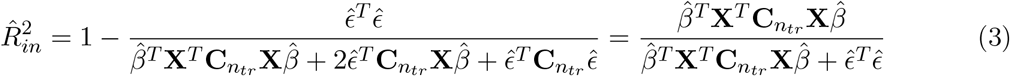

The last part of Eq. (3) matches another commonly employed definition of 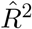 as the ratio between the variance explained by the model and the total variance in the data. Please note that this definition is identical to the one we gave in Eq. (2) *only* when using OLS and estimating the coefficient of determination in-sample. If regularization is used, or if *R*^2^ is evaluated on a different dataset, the two definitions diverge, as the orthogonality between model and residuals does not hold anymore (i.e. 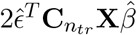 is not zero). In the remainder of the work, we will refer to estimates obtained on the training data as “*in sample*”, and the ones on the test data as “*out of sample*”.

In the next sections, we will provide an in-depth characterization of the properties of the coefficient of determination *R*^2^. We will first consider the case of OLS fit, and characterize *R*^2^ when it is computed in-sample 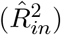 (section 2.3) and when it is computed out-of-sample 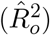 (section 2.4). We will then move to regularized regression, and illustrate its effect on *R*^2^ both in the training and testing dataset (section 2.5). Finally, we will introduce corrections for all the scenarios (in- and out-of-sample, with and without regularization) based on quadratic form approximations (section 2.6).

### 2.3 Properties of the in-sample Coefficient of Determination 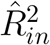

Using the hat matrix (see glossary) it is possible to map the observations *y* into their fit: *ŷ* = **H***y*, which allows the estimated residuals to be written as:

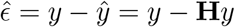

which in terms of the generative model is:

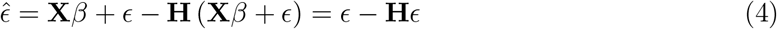

where **HX** = **X** by construction (assuming OLS is used). Eq. (4) expresses the estimated residuals as a combination of two terms: the true, irreducible error, related to the noise in the observations *ϵ*, and its OLS fit: **H***ϵ*. In the case of a large number of samples, the second term approaches zero, since **X** and *ϵ* are independent. Substituting the terms in Eq. 4 in the definition of 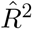, we obtain:

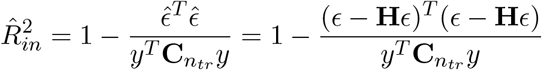

Exploiting the fact that **H** is an idempotent matrix we obtain:

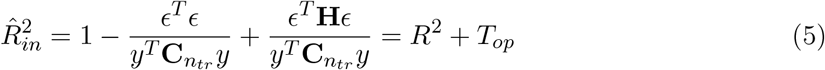

In this form, the estimated 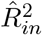 is expressed as the sum of two terms, both ratios of quadratic forms. The first, which we call *R*^2^, is not related to the fitting and is the value that can be obtained with perfect knowledge of the coefficients *β* and therefore the noise component. Here we consider this to be the *true* value of *R*^2^ *for the given sample*. This term is a random variable (the additive noise *ϵ* comes from a stochastic process) whose expected value across noise realizations and design matrices **X** can be taken as the population *R*^2^:

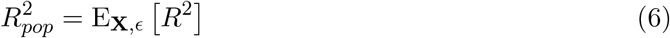

The second term, *T*_*op*_, is always non-negative and it represents the *optimistic bias* introduced by overfitting, artificially increasing the estimated 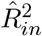 and making it a biased estimator:

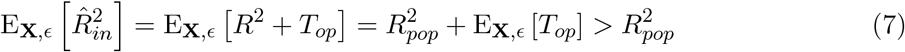

### 2.4 Properties of the out-of-sample Coefficient of Determination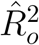

In what follows we assume that the model is fit on the training data (**X**, *y*) and evaluated on a new, independent testing dataset (**X**_*o*_, *y*_*o*_) coming from the same distribution and for which the same generative model holds, *i*.*e*.: *y*_*o*_ = **X**_*o*_*β* + *ϵ*_*o*_.

Similarly to the formulation used for 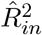 (see section 2.3), we define the Hat matrix for the out-of-sample data as **H**_*o*_ = **X**_*o*_**X**^*†*^. Please note that **X**^*†*^ is computed on the training data **X**. The predicted observations *ŷ* _*o*_ on the test data are obtained as: *ŷ* _*o*_ = **H**_*o*_*y* and the estimated out-of-sample residuals can be written in terms of the generative model as:

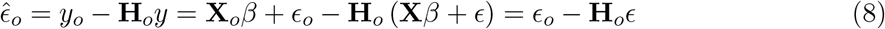

exploiting the fact that **H**_*o*_**X** = **X**_*o*_**X**^*†*^**X** = **X**_*o*_.

Equation 8 shows that there are two contributions to the estimated residuals 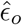: the noise in the test data *ϵ*_*o*_ and the OLS projected noise from the training data **H**_*o*_*ϵ*. This provides an intuition behind overfitting in linear regression. We showed that in the training data, the estimated residuals can be decomposed in two terms that are dependent on *ϵ* (see Eq. 4). The total variance of their combination is smaller than the error variance *σ*^2^, showing that adding predictors to the model decreases the in-sample variance. On the other hand, in the test data, the two terms in Eq. 8 are independent and therefore the variance of the estimated residuals is larger than the error variance in the test data, this phenomenon is exacerbated the more predictors are used. See [Rosset and Tibshirani, 2020] for in-depth analysis and bounds of the error of in- and out-of-sample linear regression.

Substituting the decomposition of 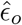 (Eq. 8) in the definition of the out-of-sample 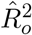 we have:

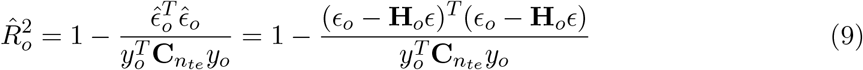

It is possible to express 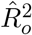 as a function of the true *R*^2^ as:

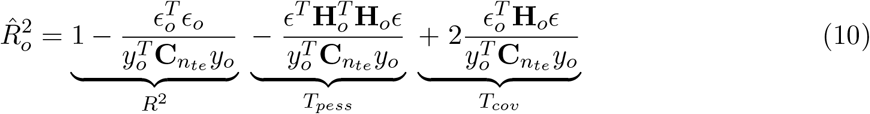

The first term, *R*^2^, represents, as in the previous section, the *R*^2^ value for the test dataset that we would obtain by assuming perfect knowledge of the additive noise *ϵ*_*o*_ (we refer here to this again as the *true R*^2^). The term *T*_*pess*_ represents the bias that fitting the noise in the training data **H**_*o*_*ϵ* has on the test data and it is always negative (hence we refer to this as pessimistic bias compared to the optimistic bias in the training data) [Rosset and Tibshirani, 2020].

It is worth noting that, when the same **X** is used in-sample and out-of-sample cases (*i*.*e*. fixed experimental design between training and test datasets), *T*_*pess*_ mirrors *T*_*op*_. This resembles the case of the error term studied in [Rosset and Tibshirani, 2020] in which the pessimism term is *identical* in magnitude and opposite in sign to the optimism term. However, in our case we are studying the fraction of explained variance, and the equality between the two terms for a fixed design matrix breaks down because of the ratio with respect to the total sum of squares, which depends on different noise terms in the training and testing data.

The third term *T*_*cov*_, is not a quadratic form and its effect on 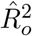 can be either positive or negative. The expected value of *T*_*cov*_ is zero since it is a function of the covariance between two i.i.d random variables (*ϵ*_*o*_ and **H**_*o*_*ϵ*).

### 2.5 Effect of regularization on the estimate of *R*^2^

The *R*^2^ properties presented in the previous sections were based on the assumption that the regression coefficients *β* are estimated with OLS. However, in most of the applications, due to collinearity or the high dimensionality of the feature spaces, i.e. (*n*_*tr*_ *<< p*), the coefficients are estimated with Ridge regression or with other variants of regularized regression. When Ridge regression is used, the estimated coefficients are shrunk (resulting in a reduction of th e scale of the estimated coefficients) and this bias has also an impact on the explained variance. The in-sample projection matrix is now: **H**(*λ*) = **X**(**X**^*T*^ **X** + *λ***I**)^−1^**X**^*T*^. In this case, the property **HX** = **X**, exploited in OLS, is not valid anymore. Consequently, the decomposition of the estimated residuals is more complex than the one shown in Eq. 4, and results in three terms:

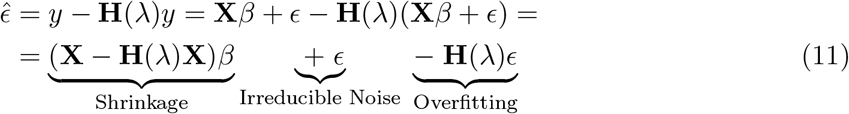

Note that, in the case of no collinearity, when *λ* = 0, the estimated error 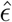 reduces to the one described in Eq. 4. Equation 11 decouples the effect of signal and noise in the context of regularization: the first term (**X** *−* **H**(*λ*)**X**)*β* corresponds to the effect of shrinkage on a noiseless dataset. This term describes the error that results in fitting a noise-free signal in the space spanned by the regularized features, rather than in the space spanned by the original features. The stronger the regularization, the more pronounced this error; in case of no regularization (therefore using OLS), this term goes to zero. The last term *−***H**(*λ*)*ϵ* is related to fitting the noise (*β* is not involved), similarly to the OLS case (eq. 4), with the difference that it is now dependent on the regularization parameter *λ*. This term decreases with increasing regularization, an opposite behavior to the shrinkage term. Selecting the optimal regularization parameter reflects finding the optimal trade-off between these two terms.

In the out-of-sample scenario we can decompose the estimated residuals 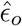 as:

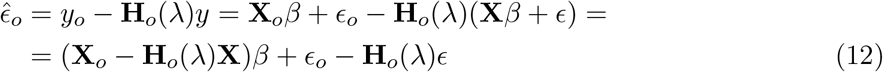

Based on these decomposition of the residuals, 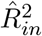 and 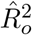 can be expressed as:

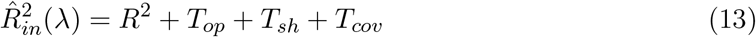

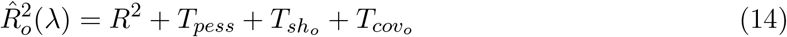

That is, compared to the OLS case, in the case of regularization 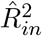 and 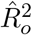 include an additional term that represents the effect of shrinkage on 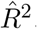. A complete description of the terms in Eq. (13)-(14) is presented in Appendix A.

The algebraic form of terms contributing to 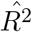 is also affected by regularization. However, their interpretation remains the same. Since **H**(*λ*) is not idempotent when regularization is used, we have:

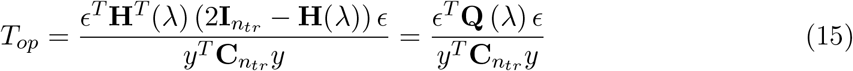

where **Q** (*λ*) = **H**^*T*^ (*λ*) (2**I**_*ntr*_ *−* **H**(*λ*)). The expression for *T*_*pess*_ is similar to the one presented in Eq. 10, with the dependency of **H**_*o*_ on *λ*.

The contribution of the shrinkage for 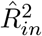 is:

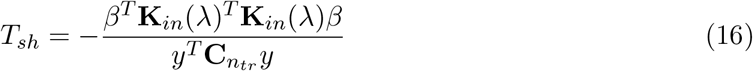

Where **K**_*in*_(*λ*) = (**I** *−* **H**(*λ*)) **X**. The matrix **K**_*in*_(*λ*) gives insights on the amount of variance of **X** that is removed as a consequence of using regularization.

For the out-of-sample *R*^2^ the corresponding term is:

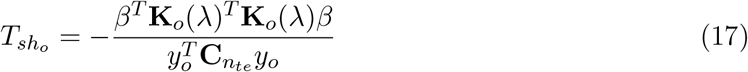

With **K**_*o*_(*λ*) = (**X**_*o*_ *−* **H**_*o*_(*λ*)**X**). A summary of these terms is presented in Table 1, second column.

**Table 1:** Bias terms in 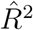, together with their approximation as a function of the true *R*^2^. In the case of OLS, *T*_*sh*_ and 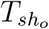 are zero. Note that the matrices **H, K** and **Q**, for both in-sample and out-of-sample depend on the regularization coefficient *λ*

| Term | Formula | Approximation | $k$ |
| --- | --- | --- | --- |
| $T_{op}$ | $\frac{\epsilon^T \mathbf{Q}(\lambda) \epsilon}{y^T \mathbf{C}_{n_{tr}} y}$ | $(1 - R^2)k_{op}$ | $k_{op} = \frac{Tr(\mathbf{Q}(\lambda))}{n_{tr}}$ |
| $T_{pess}$ | $-\frac{\epsilon^T \mathbf{H}_o^T(\lambda) \mathbf{H}_o(\lambda) \epsilon}{y_o^T \mathbf{C}_{n_{te}} y_o}$ | $(1 - R^2)k_{pess}$ | $k_{pess} = -\frac{Tr(\mathbf{H}_o^T(\lambda) \mathbf{H}_o(\lambda))}{n_{te}}$ |
| $T_{sh}$ | $-\frac{\beta^T \mathbf{K}_{in}(\lambda)^T \mathbf{K}_{in}(\lambda) \beta}{y^T \mathbf{C}_{n_{tr}} y}$ | $R^2 k_{sh}$ | $k_{sh} = -\frac{Tr(\mathbf{K}_{in}(\lambda)^T \mathbf{K}_{in}(\lambda))}{Tr(\mathbf{X}^T \mathbf{C}_{n_{tr}} \mathbf{X})}$ |
| $T_{sho}$ | $-\frac{\beta^T \mathbf{K}_o(\lambda)^T \mathbf{K}_o(\lambda) \beta}{y_o^T \mathbf{C}_{n_{te}} y_o}$ | $R^2 k_{sho}$ | $k_{sho} = -\frac{Tr(\mathbf{K}_o(\lambda)^T \mathbf{K}_o(\lambda))}{Tr(\mathbf{X}_o^T \mathbf{C}_{n_{te}} \mathbf{X}_o)}$ |

The exact estimation of *T*_*sh*_ and 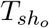 would require knowledge of the true *β*, which is by definition the goal of regression. However, both terms can be related to the true *R*^2^ via an approximation (see section 2.6). This approximation can be used to correct the effect of regularization on the explained variance. The remaining terms *T*_*cov*_ and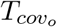, contain cross-terms between the coefficients *β, ϵ* and *ϵ*_*o*_, which are independent, and thus *T*_*cov*_ and 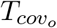 have a minor contribution to 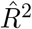 (see Appendix A).

### 2.6 Accounting for biases in 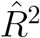

As illustrated in the previous sections, in the general case where ridge regression is used, two sources of bias affect the estimated explained variance: the bias due to overfitting (*T*_*op*_ or *T*_*pess*_) and the bias due to shrinkage (*T*_*sh*_ or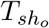). The terms *T*_*cov*_ and 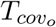 have zero contribution in expected value, and we will ignore them in the rest of this section. Due to all these terms, comparing different models using the estimated 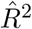, both in-sample and out-of-sample, can lead to wrong conclusions. An *unbiased* estimator of the population *R*^2^ would prevent erroneous conclusions in model comparison introduced by the different bias terms. The statistical properties of the in-sample 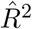 (based on OLS estimation) have been extensively studied, and several alternative estimators are available. The most common is the Ezekiel estimator [Ezekiel, 1930a], which applies a correction factor that depends on the number of features:

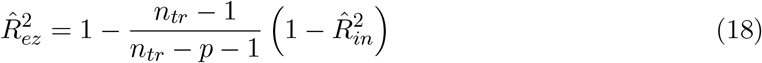

As discussed in [Karch, 2020], the Ezekiel estimator is based on the ratio of two unbiased estimates. Unfortunately, this is not an unbiased estimator of the ratio, and therefore the Ezekiel estimator does not entirely remove *T*_*op*_, meaning it is still a biased estimator (albeit the bias is much smaller when compared with 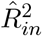). The Ezekiel estimator is not the only one proposed in the literature, see [Raju et al., 1997] and [Yin and Fan, 2001] for reviews. The “Smith correction”, in particular, described by Ezekiel in [Ezekiel, 1930b], is relevant here as it is related to the estimator we introduce:

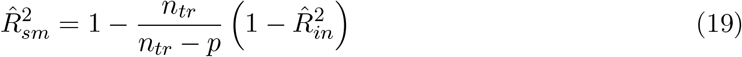

The Ezekiel and Smith estimators are asymptotically identical for increasing *n*_*tr*_. An unbiased estimator of *R*^2^ is the Olkin-Pratt estimator [Olkin and Pratt, 1958], which is computationally more demanding (a fast implementation in R has been provided by [Karch, 2020]). Another related estimator is the so-called “coefficient of cross-validation” [Yin and Fan, 2001], or “population cross-validity” [Raju et al., 1997], that estimates 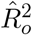 from 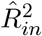 based on an analytical derivation, without the need of a test dataset for empirical evaluation. Several formulations are available, summarized in [Yin and Fan, 2001, Raju et al., 1997]. However, a good estimation of the 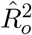 value is not useful to compare models, as we have shown that 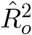 is a biased estimator of 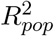.

To the best of our knowledge, no extension of the above-mentioned corrections is available in the out-of-sample scenario, or when regularization (Ridge Regression) is used. Here we develop a general adjustment based on approximating the bias sources by means of linear functions of the true *R*^2^:

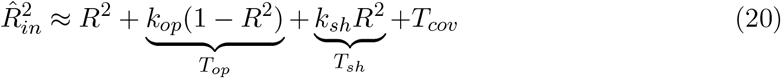

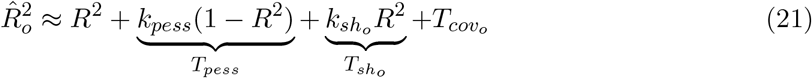

Based on these approximations and assuming that the contributions of *T*_*cov*_ and 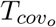 are negligible, we can obtain the adjusted 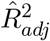 as a linear transformation of the estimated 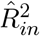 and 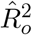. For the sake of simplicity, we will use the same notation for the in-sample and out-of-sample correction (easily disambiguated based on context):

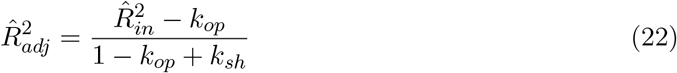

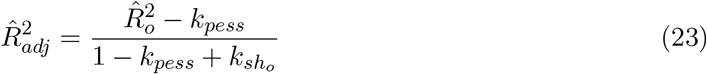

Where the constants *k*_*op*_, *k*_*pess*_, *k*_*sh*_, and 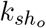 depend exclusively on the design matrices **X** and **X**_*o*_ and on the regularization parameter *λ*, not involving *y* or *y*_*o*_. Table 1 describes the approximation of each of the terms contributing to 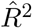, while the details of the approximations are in Appendices C and D.

It is important to note that these corrections are based on approximations and are therefore not guaranteed to result in unbiased estimates. Similarly to the Ezekiel estimator, the approximations we introduce are based on estimating the expected value of a ratio by taking the ratio of the expected values of the numerator and denominator (see Appendix B for the approximation of quadratic forms) (see [Karch, 2020]). In the out-of-sample case, the numerator and denominators are based on two independent noise terms (*ϵ* and *ϵ*_*o*_), therefore the approximation holds better. The terms *k*_*op*_ and *k*_*pess*_ are always present, while the shrinkage terms *k*_*sh*_ and 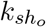 are zero when OLS (*λ* = 0) is used.

In the case of correcting 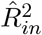 when OLS is used, the hat matrix **H** is idempotent and the analytical expression of *T*_*op*_ falls back to Eq. (5) and the numerator of the term *k*_*op*_ simplifies to *Tr* (**H**) = *p*, if **X** is non-singular. Therefore, 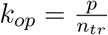, which results in the Smith estimator described above. In fact, considering Eq. 19 and expressing 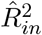 as a function of *R*^2^, we obtain:

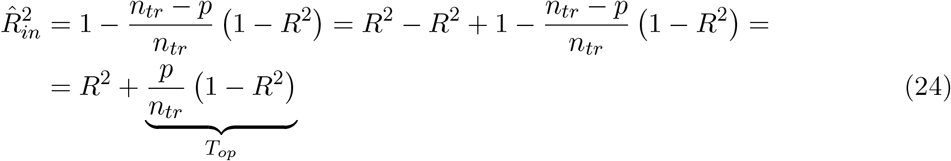

In the out-of-sample case, *k*_*pess*_ cannot be simplified, but the terms presented in Table 1 can be used for adjusting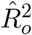.

### 2.7 Simulations

We introduce a generative model to simulate data for the numerical evaluation of the introduced approximations. Feature spaces **X**_**1**_ and **X**_**2**_ were generated from a partitioned multivariate normal distribution:

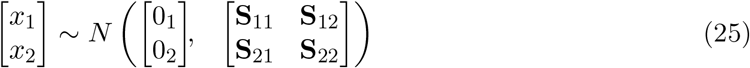

and the observed responses were generated as follows:

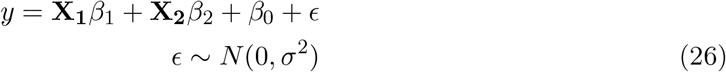

where *β*_0_ is the intercept. While the feature matrices **X**_**1**_ and **X**_**2**_ are assumed random across repetitions of the same simulation, the regression coefficients *β*_1_ and *β*_2_ and the scalar *β*_0_ are assumed fixed across the simulations. This generative model allows to generate data for model comparison, as well as, for studying *R*^2^ corrections in a single feature space (*X*_2_ = *∅*). Note that this is the generative model that is *assumed* when variance contributions of different feature spaces are estimated using *commonality analysis* [Seibold and McPhee, 1979]. We consider the simpler case of **S**_12_ = 0. However, this does not compromise the generalization of our results to the more general case where there is non-null block covariance between feature spaces (shared component), see Section 3.2. Under this generative model, the population explained variance 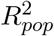 for each feature space, is computed as the average across all generated datasets of the *R*^2^ computed with perfect knowledge of the model coefficients (see Eq 5).

### 2.8 Imaging data and analyses

We used the publicly available dataset from the Algonauts 2023 challenge [Gifford et al., 2023], a competition that pursues the prediction of brain response to natural scenes in areas of the visual pathway. The data is part of the NSD dataset [Allen et al., 2022], a large-scale fMRI measurement of the brain responses to 10000 natural scene photographs in eight subjects. Each image was presented three times and the average brain response of each voxel in the visual cortex is provided. We used the training data from S01 (subject one), consisting of the brain responses to the presentation of 9841 images. We only focused on the training data, since the brain responses in the test data were not provided to the challenge participants. Additionally, for the sake of illustrative purposes, we restricted our analysis to left-V1 ROI which contains 710 voxels.

We used a linear encoding approach, where the brain responses of each voxel were modeled as linear functions of a set of pre-defined features. We arbitrarily selected, as feature sets, the ‘fc8’ and the ‘fc6’ layers from AlexNet [Krizhevsky et al., 2012], containing *p*_*fc*8_ = 1000 and *p*_*fc*6_ = 4096 activations respectively. We compared these feature spaces in their ability to explain the observed brain responses. The available data was randomly split into a training (8841 images) and a testing dataset (1000 images). To keep the analyses consistent with our simulations, we modulated the sample size by constructing different datasets from the training dataset, by starting from a random selection of 500 images and adding at random images until covering the 8000 images. Ridge regression with a fixed lambda (*λ* = 10^2^) was used to compute the linear encoding coefficients. The in-sample and out-sample *R*^2^ were computed as well as the corresponding adjusted values.

## 3 Results

### 3.1 Simulations

We consider the *n*_*tr*_ *> p* scenario in which 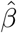 is estimated with OLS, and the *n*_*tr*_ *< p* scenario in which 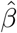 is estimated with Ridge regression. We evaluate the effect of the training set size, the model dimensionality, the test set size, and the amount of regularization separately (while keeping all other parameters fixed). Each simulation was repeated 1000 times, and the mean and confidence interval of the estimators was obtained by bootstrap across simulation repetitions. The MATLAB [MathWorks, 2020] source codes to reproduce the simulations, the real data analysis, the figures, and the implementation of the 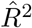 adjustment are available at https://github.com/mlsttin/adjustingR2. The function to perform the 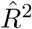 adjustment is also available in R and Python.

#### 3.1.1 Numerical evaluation of the in-sample 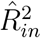

Figure 2, top panels, presents the simulation results for 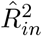 and 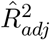, in the *n*_*tr*_ *> p* case. We present the influence of the training set size on the terms contributing to 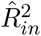 (Figure 2 top left), see Eq. 5. The number of features was fixed to *p* = 10. As expected, 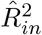 shows a positive bias that decreases with the amount of training data, suggesting convergence to the population value (green dashed line) as the number of training samples increases. Therefore, 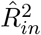 is a consistent estimator, albeit a biased one. The adjusted explained variance 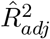 (purple curve) shows instead no (or undetectable) bias across the explored range of training set sizes.

**Figure 2:**
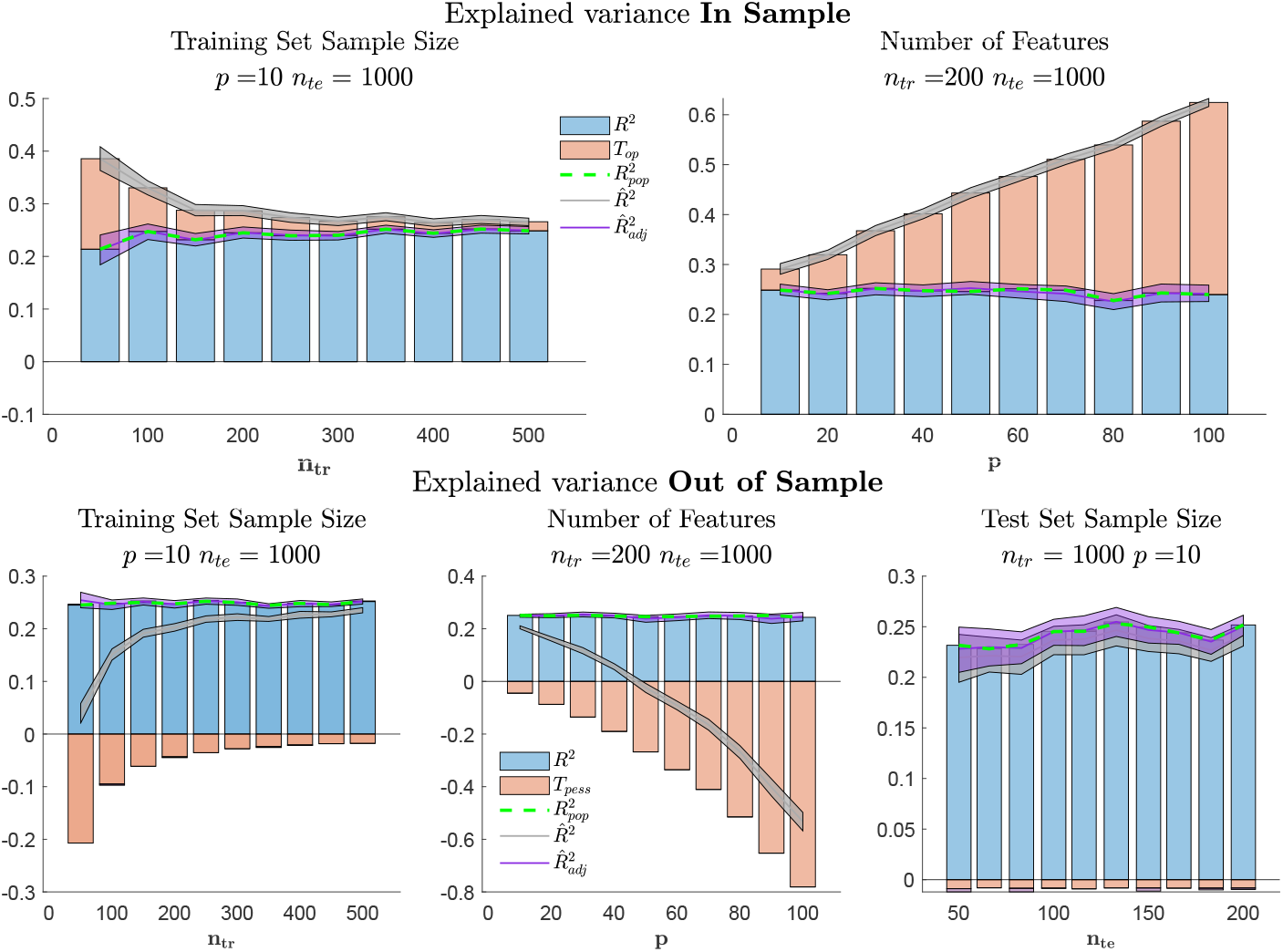
Top panels: terms contributing to 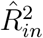 as a function of the training set size (left), model dimensionality (right). Bottom panels: terms contributing to 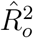 as function of training set size (left), model dimensionality (central) and test set size (right). The gray line and the purple line describe the unadjusted and adjusted *R*^2^ estimators respectively. The model coefficients were estimated with OLS, the shadow areas describe the 95% confidence intervals of the mean, obtained with bootstrap. The population *R*^2^ is described by the green dashed line.

Analyzing the contribution of *T*_*op*_ to the bias of 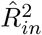 confirms the results obtained analytically in the previous sections. That is, the optimistic bias of 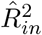 is determined by *T*_*op*_, a term related with noise fitting. The value of *T*_*op*_ approaches zero for large training samples, a consequence of the independence between **X** and *ϵ*.

Figure 2 top right panel presents the effect of model dimensionality on 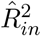 and 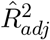 (with the training data size fixed to *n*_*tr*_ = 200). Confirming well established results in linear regression, the bias in 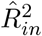 linearly increases with the number of features in the model, while 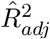 corrects for this bias and, independently of the model dimensionality, approximates the population value (green dashed line). Also in this case, the bias of 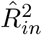 is determined by *T*_*op*_, highlighting the increased contribution of overfitting as model dimensionality increases.

#### 3.1.2 Numerical evaluation of the out-of-sample 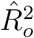

Figure 2, bottom panels, reports the simulation results for the estimated out-of-sample explained variance 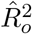. In the left panel we present its dependence on the training set size (model dimensionality is fixed to *p* = 10). The number of testing samples was fixed to a large value to discard its influence on the results (*n*_*te*_ = 1000). 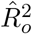 is biased in the opposite direction compared with 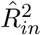. This bias decreases with the amount of training data. The bias in 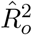 is explained by *T*_*pess*_, see Eq. (10), which reflects the effect of overfitting the training dataset on the performance on the test data. *T*_*cov*_, which represents the covariance between the (rotated) training error and the test error, has a contribution to 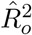 of two orders of magnitude smaller than *T*_*pess*_ and cannot be therefore appreciated from the figure. Confirming the analytical deductions of Eq. (23), the observed bias is corrected by 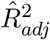.

In the central panel we report the effect of model dimensionality on 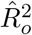 (with fixed training sample size and test size: *n*_*tr*_ = 200, *n*_*te*_ = 1000 respectively). The negative bias of 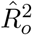 increases with the number of features in the model. It is important to note that in this case, the effect is non-linear. Also, in this case, the contribution of 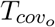 is negligible. The results indicate that the introduced 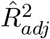 corrected the 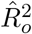 bias across the explored range of model dimensionalities.

In the right panel, we illustrate the effect of changing the amount of test samples on the estimation of 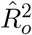. In this scenario, we considered *p* = 10 and *n*_*tr*_ = 1000 to reduce the effect of overfitting the training data. The results indicate that the amount of test data has a smaller impact on the bias in the observed 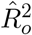, and that the proposed adjustment correctly recovers the population *R*^2^ value.

We have shown that 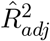 is an unbiased estimator of the population *R*^2^, in that it corrects the bias induced by the two terms *T*_*op*_ and *T*_*pess*_. As discussed in section 2.3, the true *R*^2^ is specific to the noise realization in a given dataset, and it is equal, in expected value, to the population value 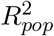. It is therefore interesting to relate the variability across datasets of 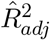 to the variability of the true *R*^2^. In Figure 3 we show both the in-sample estimator, the out-of-sample estimator, and the adjusted 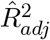, versus the true *R*^2^ across the datasets with the same simulation setting. For the in-sample 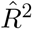 (left panel) we observe that the variability of 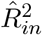 and 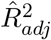 is mainly due to the variability of the true *R*^2^, which is due to variations in **X** and *ϵ* (under the same *β* in the generative model). For the out-of-sample 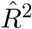 we observe that the increase in the amount of test data (*n*_*te*_ = 50 central panel, up to *n*_*te*_ = 1000 right panel) has a modest effect on the reduction of the variance of 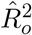 and 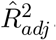. This can be explained observing that the variance of 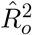 is the sum of two terms: the variance of the true *R*^2^, which depends only on the test data, and the variance of *T*_*pess*_, which depends also on the training data (see Eq.(10)). By increasing the amount of test data we reduce the variance of the first component, while the second component is mostly unaffected.

**Figure 3:**
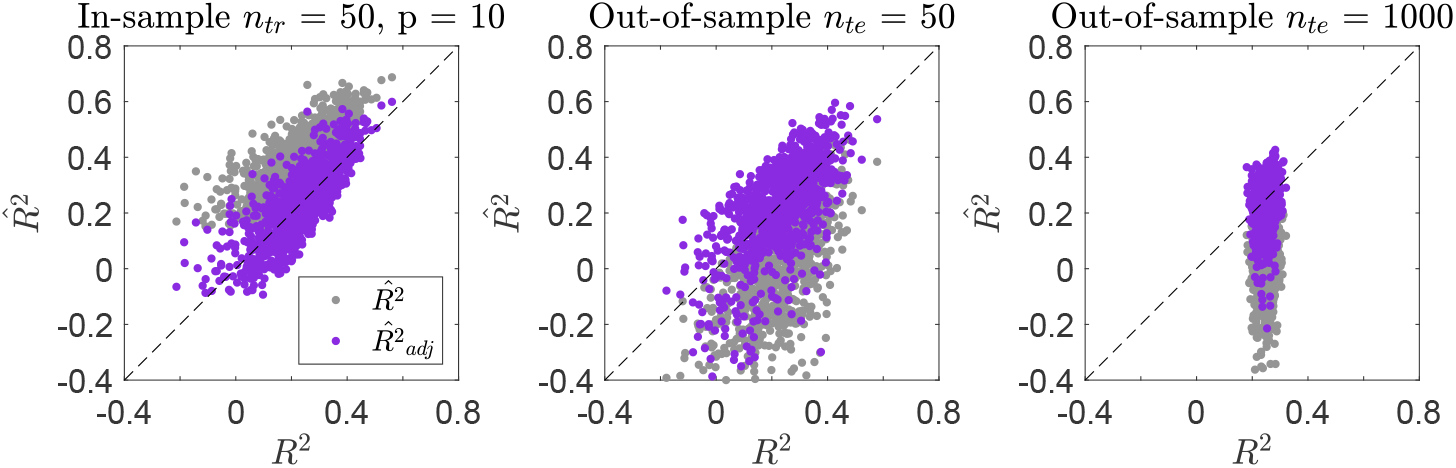
Scatter plot between true *R*^2^ and the estimated 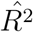 for in-sample (left), out-of-sample *n*_*te*_ = 50 (center) and out-of-sample *n*_*te*_ = 1000 (right). The gray dots and the purple dots denote the corresponding values of the unadjusted and adjusted *R*^2^ respectively. Each point in the figure corresponds to one simulation.

#### 3.1.3 Numerical evaluation of 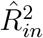 and 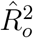 with regularization

In Figure 4 we present each of the contributing terms to 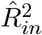 and 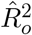, together with the proposed correction, and the population value 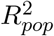, in the *n*_*tr*_ *< p* case. We examined the effect of the number of training samples (fixing the model dimensionality to *p* = 100 features) and the effect of the model dimensionality (fixing *n*_*tr*_ = 50 samples). In all cases, we used Ridge Regression and fixed the regularization parameter to *λ* = 100.

**Figure 4:**
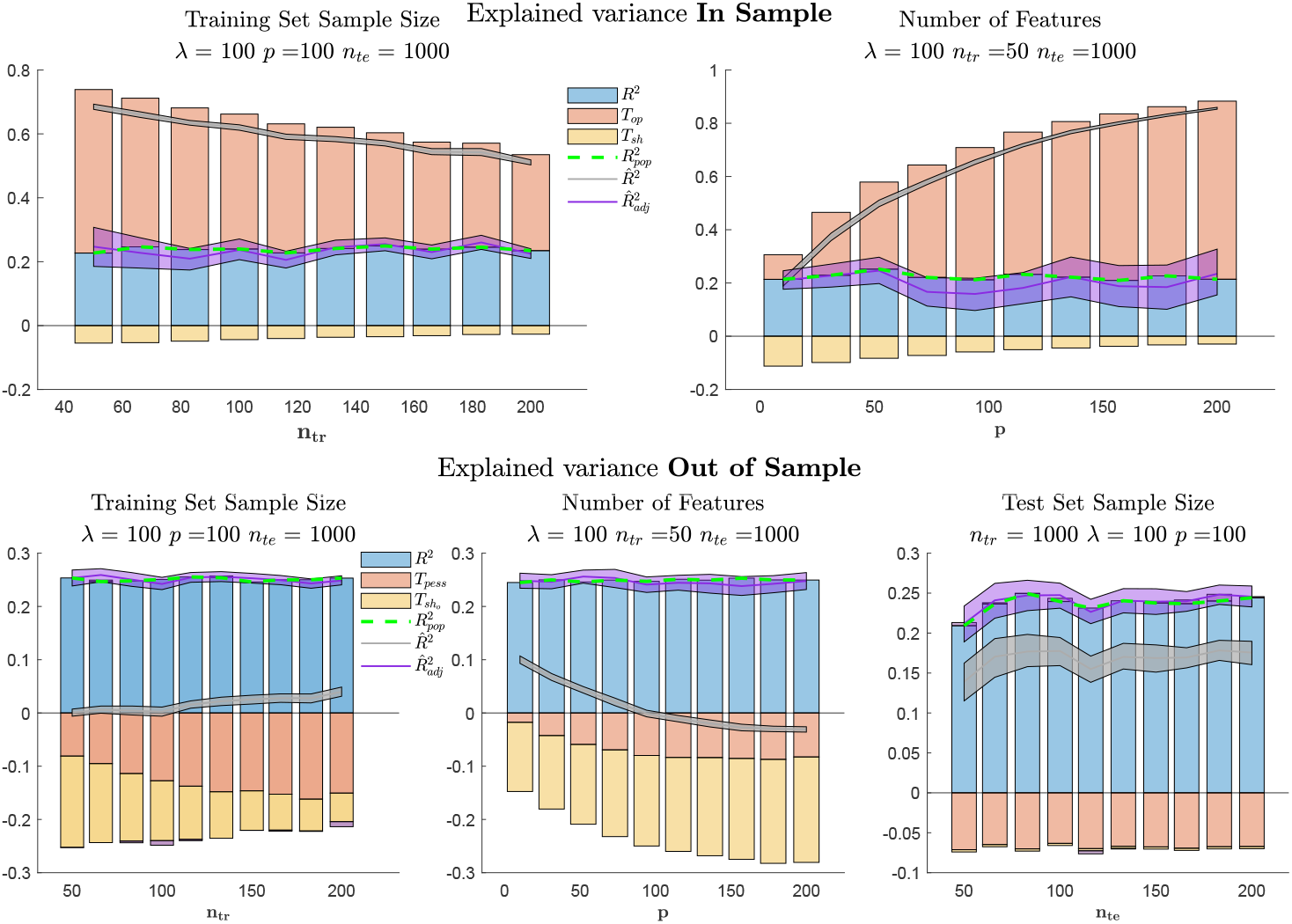
Top panels: terms contributing to 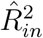 as a function of training set size (left), model dimensionality (right). Bottom panels: terms contributing to 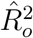 as a function of training set size (left), model dimensionality (central) and test set size (right). The gray line and the purple line describe the unadjusted and adjusted *R*^2^ estimators respectively. The model coefficients were estimated with Ridge regression with *λ* = 100. The population *R*^2^ is described by the green dashed line.

In the top panels, we focus on 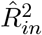. As discussed in section 2.3, 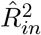 is the result of a trade-off between *T*_*op*_ and *T*_*sh*_. These two terms have opposite contributions to the estimated explained variance. The proposed correction, 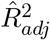, (purple line in Figure 4) accounts for both sources of bias, and approximates 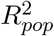 (green dashed line).

The bottom panels report the results for the out-of-sample estimator. In this case, *T*_*pess*_ and 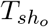 contribute to the observed 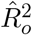in the same direction, reducing its value. Also in this case, the 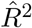 estimated using the proposed approach (purple line) corrects both sources of bias. Interestingly, in this simulation, the adjusted estimator showed less variance compared to the in-sample case. For a small training set size (bottom-left panel), the effect of shrinkage is larger, and it becomes less relevant the larger the training size (for a fixed *λ*). In the bottom-right panel, we show the effect of changing the test sample size *n*_*te*_ (fixing the training size to *n*_*tr*_ = 1000 and model dimensionality to *p* = 100). These results suggest that, similarly to the OLS case, the amount of test data has a smaller influence on the observed and adjusted 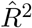 than the amount of training data.

We report the effect of model dimensionality in the top right (in-sample) and bottom center (out-of-sample) panels of Figure 4 (with *n*_*tr*_ = 50, and *λ* = 100). Compared to the non-regularized case (see Figure 2), regularization reduces the sensitivity of both *T*_*op*_ and *T*_*pess*_ to the model dimensionality. The effect of shrinkage (*T*_*sh*_ and 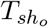) is more notable when *n*_*tr*_ *< p* (note that *λ* was fixed to a constant value in these simulations). In both in-sample and out-of-sample, 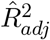 (purple line) reduces the bias in the estimated *R*^2^, with a smaller variance for the out-of-sample estimator.

The effect of the regularization parameter is presented in Figure 5. For 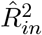 (left panel), the amount of optimism (overfitting) reduces with the amount of regularization, while the contribution of shrinkage follows the opposite pattern. Consistently, for 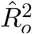 the amount of pessimism reduces with the amount of regularization, while the effect of shrinkage increases. For small to moderate regularization, the adjusted estimator 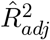 corrects the bias in both estimators 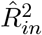 and 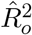, with less variance in the adjusted 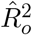. However, for large values of *λ* the 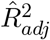 failed to correct the observed 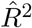 bias. This occurs because the 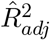 is based on a linearization that fails for large *λ* values. When substantial shrinkage is applied, numerical instabilities are induced by the ratio of small quantities in equations (43)-(44).

**Figure 5:**
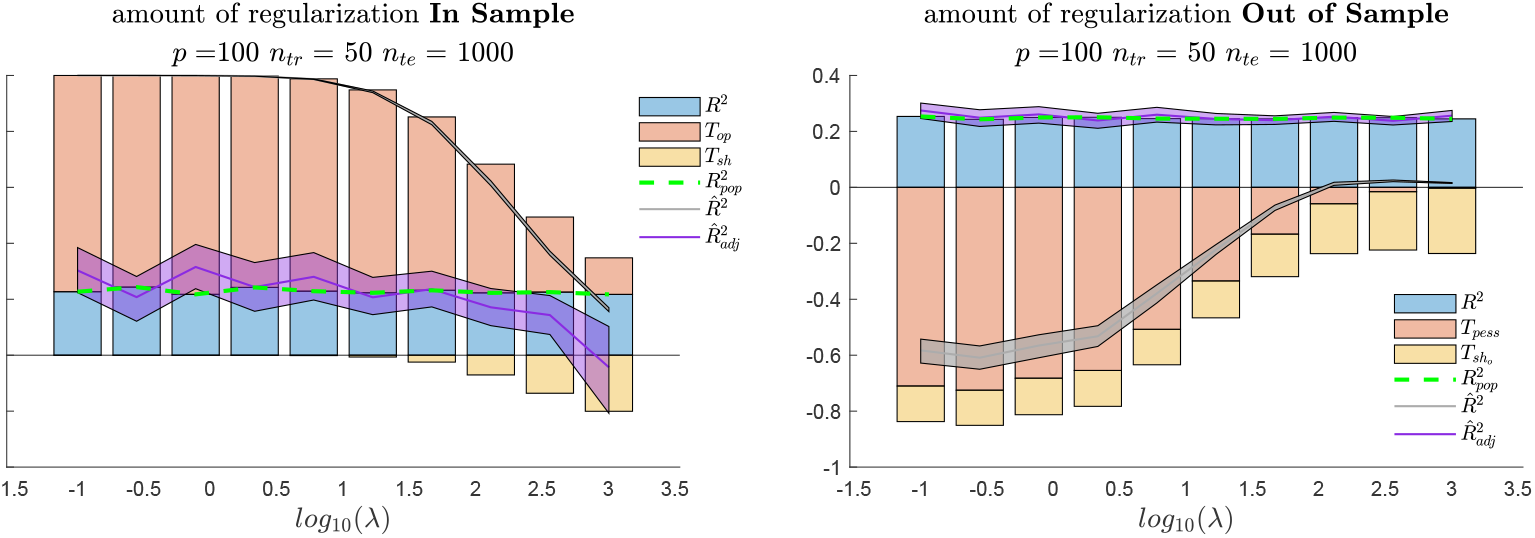
Effect of the amount of regularization *λ* on the terms contributing to 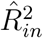 (left) and 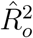 (right). The uncorrected and corrected estimators are presented in gray and purple respectively. The shaded area denotes the 95% percentile bootstrap confidence intervals. The population *R*^2^ is described by the green dashed line.

In Figure 6 we show, similarly to Figure 3, the variability of the estimated and adjusted 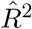 versus the true *R*^2^, for a given regularization value. The left panel describes the in-sample case. We chose the most difficult situation (*p* = 2*n*_*tr*_) to illustrate the trade-off between variance and bias in the 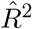 correction. That is, whereas 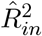 is heavily biased towards larger values with reduced variance, the correction recovers the true value, albeit with larger variance. In the center and right panels we compare true and estimated *R*^2^ in the out-of-sample case (with *n*_*te*_ = 50 in the center panel and *n*_*te*_ = 1000 in the right panel), for the same training setup as in the left panel (*p* = 100, *n*_*tr*_ = 50). The plots indicate that the bias in the 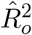 is less pronounced on the test data (albeit with larger variance), and the variances of both 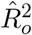 and 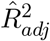 are comparable, suggesting that the adjusted out-of-sample estimator should be preferred when regularization is needed.

**Figure 6:**
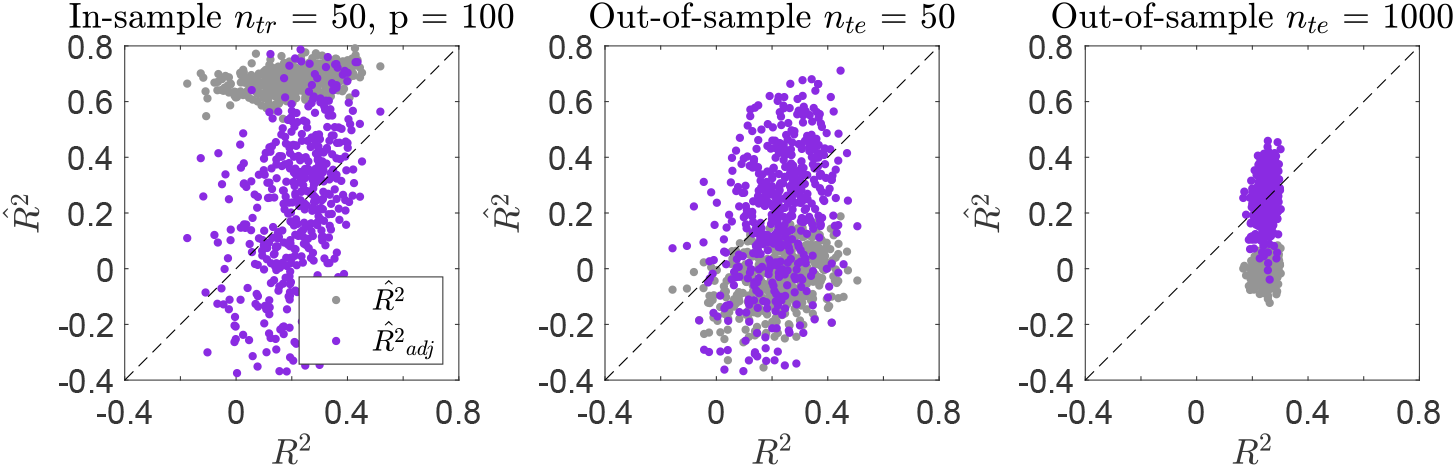
Scatter plot between true *R*^2^ and the estimated 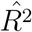 for in-sample (left), out-of-sample *n*_*te*_ = 50 (center) and out-of-sample *n*_*te*_ = 1000 (right). The gray dots and the purple dots denote the corresponding values of the unadjusted and adjusted *R*^2^ respectively. Each point in the figure corresponds to one simulation, under the setting specified in the figure title.

### 3.2 Implications for model comparison

We consider the effect that the bias in 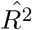 has on model comparison. We illustrate these effects with a simulation in which we compare a smaller feature space with *p*_1_ = 10 dimensions, explaining 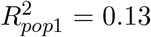 of the variance of *y*, with a larger feature space with *p*_2_ = 20 dimensions that explains 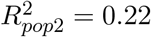. For each feature space, we estimated the regression coefficients with OLS separately and compared the obtained 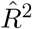. Please note that this is equivalent to first performing commonality analysis on the two models [Seibold and McPhee, 1979], and then comparing the uniquely explained component of each model. In fact, in commonality analysis the unique variance explained by each of the two models is estimated as:

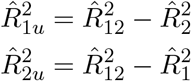

with 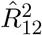 being the *R*^2^ estimated using both features spaces, and 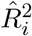 being the *R*^2^ estimated using only feature space *i*. Therefore, the difference in explained variance by the unique components 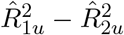, is equivalent to the difference in the estimated *R*^2^ using the two models separately 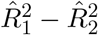.

The unadjusted and adjusted 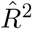 were computed for increasing values of training set sizes, for the out-of-sample we used *n*_*te*_ = 1000. For each training set size 1000 datasets, under the same simulation setting were created. For the in-sample estimators (Figure 7, left column) we observed the largest positive bias in the model with the largest dimensions *p*_2_ = 20 (second row). This is reflected by the largest negative bias in the out-of-sample 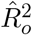 (right column, second row). The adjusted 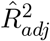 estimates corrected those biases for the in-sample and out-of-sample estimators for both feature spaces (see purple lines). The bottom row shows the differences between the explained variances of each feature space. For the in-sample estimators (left column), the unadjusted differences successfully determined that the largest model (M2) explains more variance than the smaller (M1). However, the magnitude of the difference was overestimated for small *n*_*tr*_. Importantly, for the out-of-sample estimators, the unadjusted differences produce, for small *n*_*tr*_, an incorrect conclusion regarding which of the two feature spaces explained more variance (i.e the unadjusted difference is positive, suggesting that M1 explains more variance than M2). This misleading conclusion is a consequence of the different convergence rates to the population 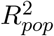 (green dashed lined) of feature spaces with different numbers of dimensions. The adjusted differences 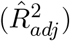, instead, correctly identified the feature space that explains more variance for the in-sample and the out-of-sample estimators, and also the magnitude of the difference.

**Figure 7:**
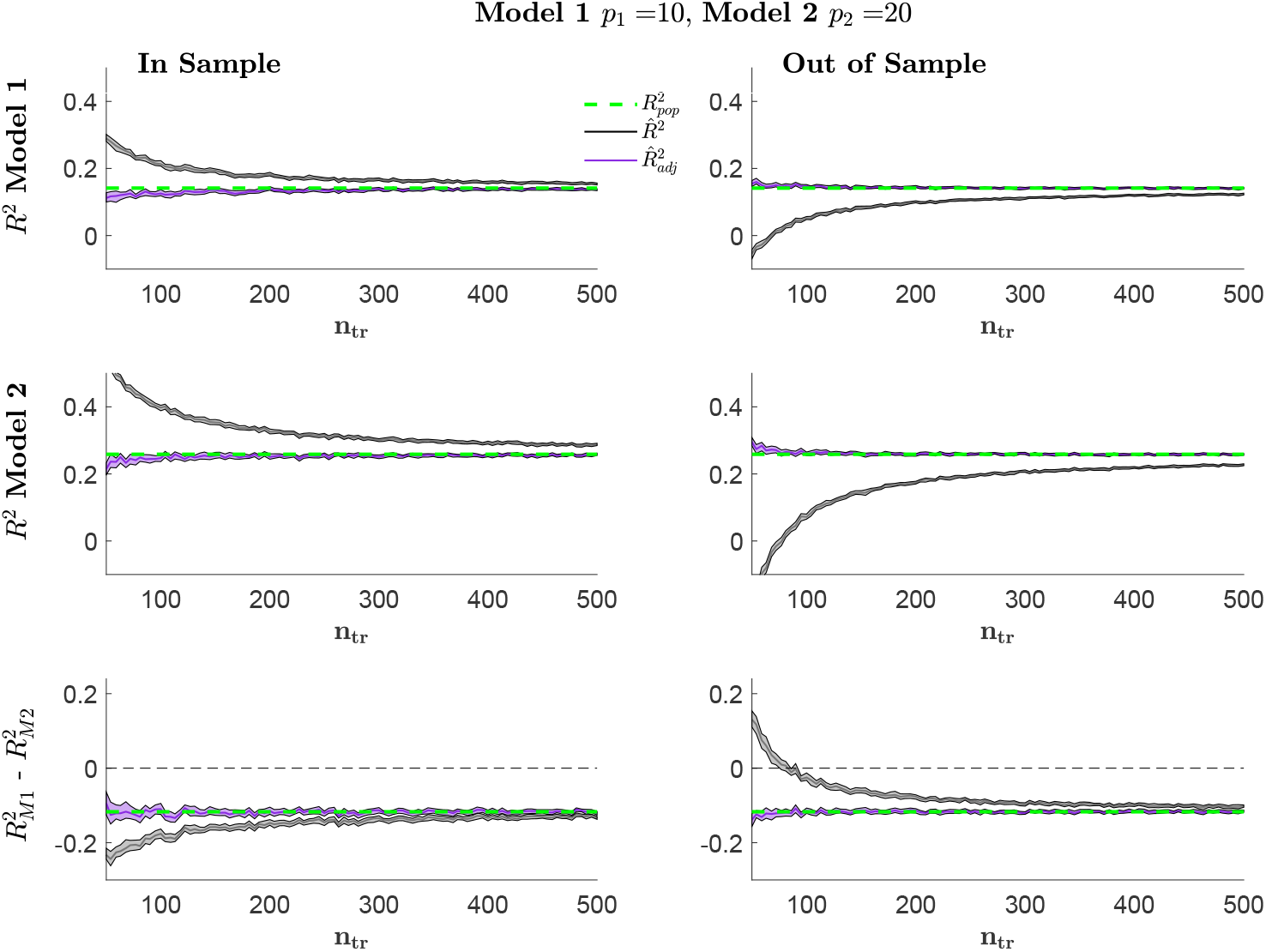
The top panel presents the 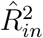 (gray) and 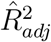 (purple) for the in-sample (left) and the out-of-sample (right) as a function of the training set size for the model of smaller dimensionality *p*_1_ = 10. The central panel presents the corresponding figure for the model of largest dimensionality *p*_2_ = 20. The bottom panel presents the unadjusted and adjusted differences between *R*^2^ of the smallest and the largest model. The green line depicts the population 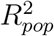 for each model and the population difference between models in *R*^2^ in the bottom panel. The shadow areas denote the 95% confidence intervals of the mean obtained with bootstrap.

### 3.3 Imaging data

#### 3.3.1 Imaging Data

Figure 8 presents the estimated explained variance obtained from a linearized encoding model, mapping the last fully connected layer of AlexNet [Krizhevsky et al., 2012], with *p*_*fc*8_ = 1000 features to the observed (fMRI) brain responses in area V1 (710 voxels). By tuning the number of training samples used, we encompassed both the *n*_*tr*_ *< p* and the *n*_*tr*_ *> p* scenario. In all cases, we used Ridge Regression, *λ* = 100 and the test set was fixed to *n*_*te*_ = 1000. To create a benchmark of the explained variance in the context of a large sample, we compute 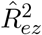 (see Eq. (18)) obtained with OLS on the whole dataset (*n*_*tr*_ = 9841 images, both training and testing). We will refer to this value as 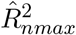. As discussed in section 2.6, the Ezekiel estimator has a relatively small bias, and we can consider 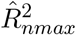 as an estimate of the 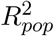.

**Figure 8:**
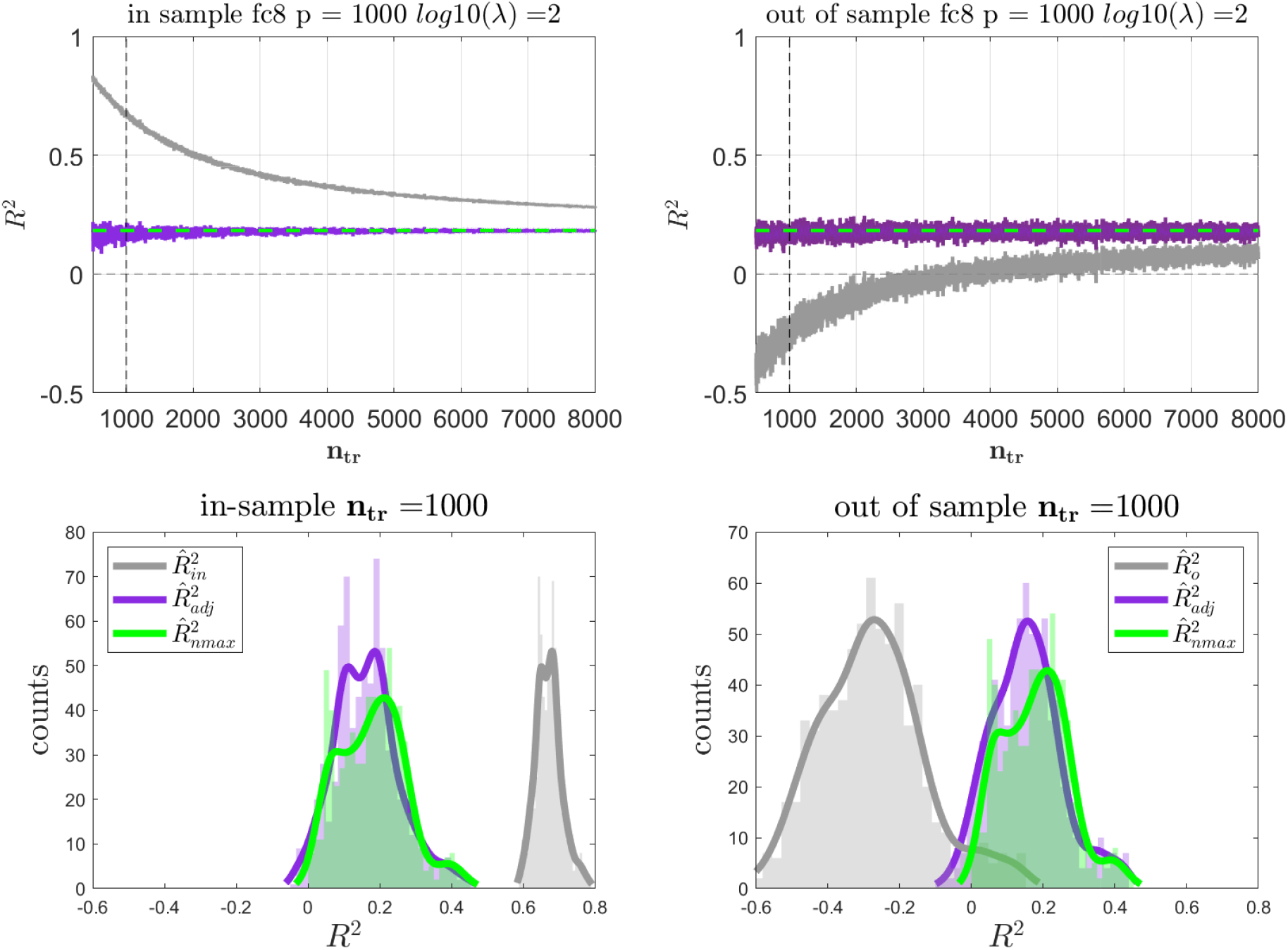
The top panel presents the estimated *R*^2^ values as a function of the training set size for the *fc*8 feature space for the in-sample (left) and out-of-sample (right) estimators. The estimated 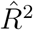 is presented in gray, the adjusted 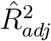 is presented in purple, while the large sample 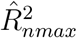 is presented in green. On the bottom panels, the distributions for the estimators are shown for an arbitrarily selected training sample size of *n*_*tr*_ = 1000.

In Figure 8, top left panel, we present 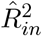, when the amount of training data monotonically increased from 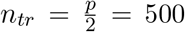 up to *n*_*tr*_ = 8*p* = 8000. The unadjusted 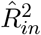 produced optimistically biased values (gray line), and the amount of optimism decreased with the increase of training data *n*_*tr*_. Consistently with the simulations, the adjusted estimate 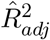 (see Eq. 23) recovers the large sample 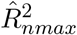. However, the correction exhibits a larger variance for small *n*_*tr*_. On the right panel, we show the corresponding figure for the out-of-sample 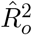. The optimistic bias of 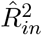 is reflected in a pessimistic bias of 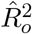, which now approaches the large sample 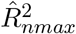 from below (even showing negative values). This bias is satisfactorily corrected by 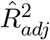. It is necessary to emphasize that the validity of the proposed correction is also a function of *λ* and tends to be less precise when larger regularization values are employed.

In Figure 8, bottom panels, we focus on one training set size set (*n*_*tr*_ = *p* = 1000, dashed vertical lines in the top panels) and present the distribution of 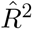 (across all the voxels, in-sample on the left and out-of-sample on the right). The adjusted 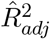 overlaps with the large-sample 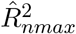, while 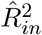 exhibits a positive bias (left panel) and 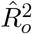 a negative bias (right panel). Importantly, these results show that fitting the model with *n*_*tr*_ = 1000 samples and applying the proposed adjustment produces 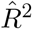 values highly comparable to those obtained when fitting the model with a training set size of *n*_*tr*_ = 9841 samples.

#### 3.3.2 Comparison between models

In the previous section, we demonstrated the performance of the *R*^2^ estimators when one layer from AlexNet (*fc*8) was mapped to the brain response in the area V1. In this section, we considered the sixth fully connected layer *fc*6, and the layer *fc*8, and compared them in terms of the variance they explained in the observed fMRI responses. Importantly, the last fully connected layer consists of *p*_*fc*8_ = 1000 features, while the sixth fully connected layer consists of *p*_*fc*6_ = 4006 features. Since the latter model is larger, we expect that model comparison based on the 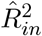 will favor the larger model (*fc*6), while an analysis based on 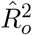 will favor the smallest model (*fc*8). The selection of these two layers was motivated by the need to have a reference value (a proxy of 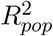 in the simulations) against which we could compare 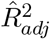. Since the dimensionality of both models was smaller than the maximum amount of training data available, we used the difference between the model’s 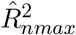 as the proxy for the ground truth (see previous section). We expect that the adjusted in- and out-of-sample estimators should ideally approach this value.

The same settings presented in the previous section were used in this analysis. In Figure 9 top panel, we present the difference 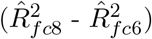, for the in-sample (left) and out-of-sample (right). When comparing the models on the training data (in-sample), *fc*6 appears to explain a larger amount of variance (negative 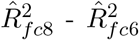, gray line). While this comparison allows correctly selecting the best model (i.e. the sign of 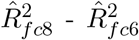 is similar to the one of 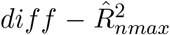, green line), the magnitude of the difference is inflated. This inflation is a consequence of the largest amount of optimism in the model with a larger number of features (*fc*6). The difference between the corrected estimators (purple line) is, on the other hand, close to the large sample difference 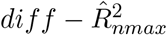.

**Figure 9:**
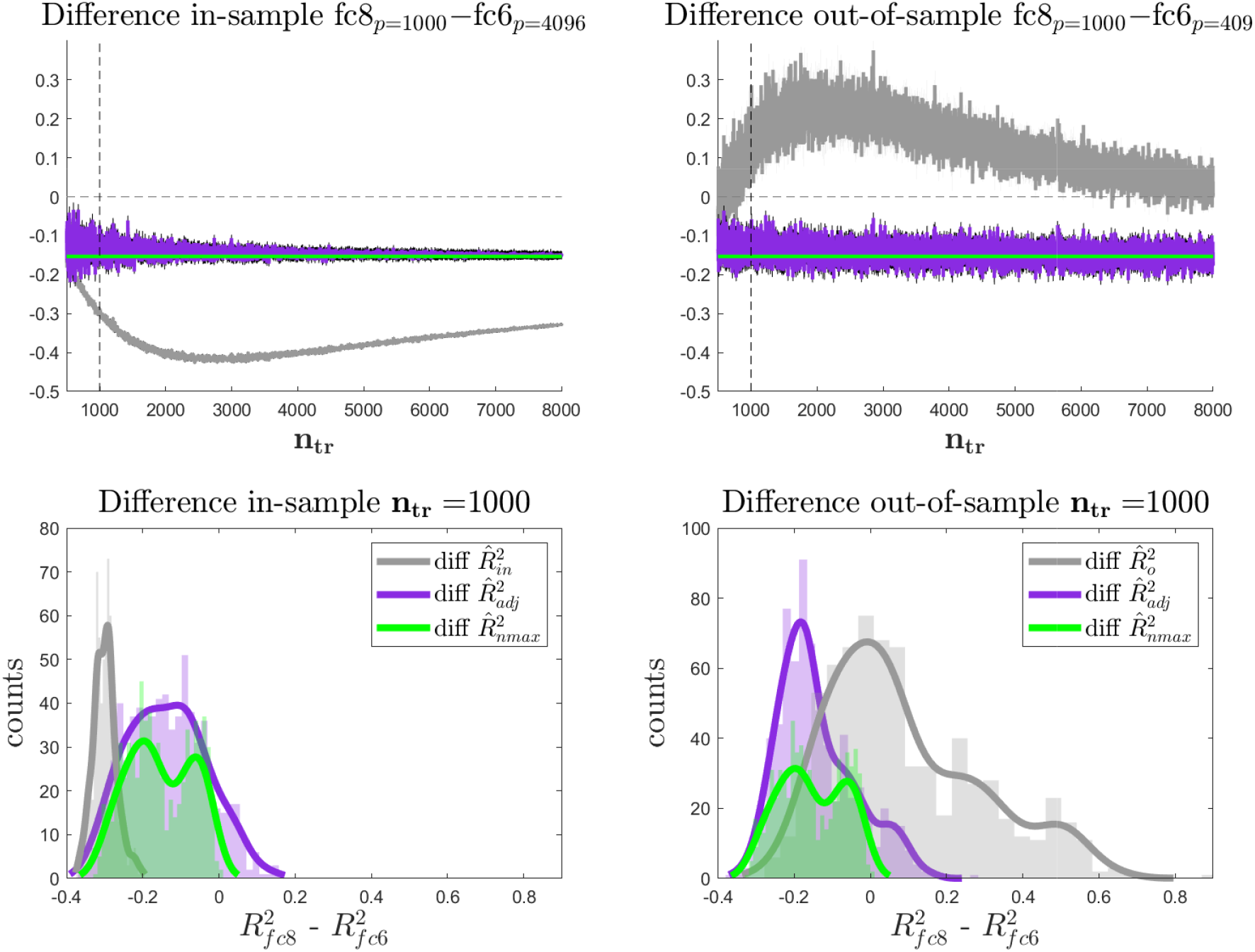
On the top panel, the difference in variance explained between feature space *fc*8 and feature space *fc*6 is presented for the in-sample (left) and out-of-sample (right) estimators. The difference between the estimated 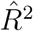 is presented in gray, the difference between the adjusted 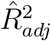 is presented in purple, while the difference obtained using OLS and a larger data sample 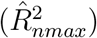 is presented in green. In the bottom panels, the distribution of the difference is presented for an arbitrarily selected training sample size of *n*_*tr*_ = 1000

The difference between uncorrected 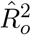 is positive across most of the range of training sample sizes (right panel). That is, using the unadjusted 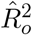, the model with smaller dimensions *fc*8 explained more variance in the test data. This conclusion, which is misleading if compared to the 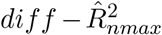, is a consequence of the different amount of overfitting for the two models (for fixed *n*_*tr*_). Importantly, using the proposed correction, 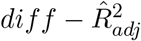 (purple line), allowed us to recover both the sign and the magnitude of the population differences in *R*^2^.

## 4 Discussion

In this work, we investigate the statistical properties of the coefficient of determination *R*^2^ in the context of multiple linear regression, characterizing both its in-sample and out-of-sample estimation. Through an explicit analysis of the underlying generative model, we describe the effect that the number of features and the available number of samples have on the bias terms affecting the *R*^2^ estimation, relating this to *overfitting* the noise component of the training data. Our theoretical approach allows us to derive a unifying framework describing the properties of the *R*^2^ for OLS as well as in the case of regularization.

In the machine learning literature, the out-of-sample model performance is used as an indicator of a model’s ability to generalize to new data. However, in fields like neuroimaging, machine learning methods are often employed not solely for prediction, but also for inferential purposes. In such instances, the primary concern shifts from assessing model performance to the inference of population parameters, which are then utilized for subsequent interpretations.

In this article, we analytically show for the coefficient of determination, how the amount of overfitting on the training data (captured by *T*_*op*_) is reflected by a drop in performance on the test data *T*_*pess*_. Making a more abstract parallel with a classifier, a model with no overfitting would generalize (both in training and in testing data) at the Bayes error rate (*i*.*e*. the error that would be obtained if perfect knowledge of the distribution of the classes was available). In our formulation, this is the 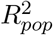, which can be obtained with perfect knowledge of the generative model, and depends only on the noise distribution.

The bias present both in the in-sample and out-of-sample estimators of the coefficient of determination may not represent a problem when the performance of a model is compared against the null distribution (i.e. when trying to reject the null hypothesis of the model having no relation with the data). A comparison of the values of 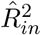 or 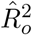 with the null distribution obtained, for instance, with a permutation or randomization test [Valente et al., 2021], would result in a valid conclusion, as both the non-resampled and resampled values are affected by the same bias. However, when the question is the one to compare the explained variance of two models (an approach that is becoming increasingly relevant when e.g. testing computational models of brain processing using linearized encoding), the bias terms can have a dramatic effect if the dimensions of the models are different. This is the case because, as we show here, the bias in 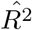 depends on the model dimensionality and different dimensionality lead to different biases and thus potentially erroneous conclusions (see e.g. Figure 1 and Figure 9). It is therefore of paramount importance to use *unbiased* estimators of 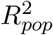 when two models are compared in terms of explained variance. It is noteworthy to highlight that, contrary to the case of fitting a single model, a permutation test based on shuffling the independent variable *y* would not help in reaching a correct conclusion on model comparison. This is because, such permutation provides samples of the null distribution testing the hypothesis of both models not explaining the data, rather than testing the null hypothesis of the two models explaining the same variance. To the best of our knowledge, a permutation test allowing to determine the null distribution of two models explaining the same variance is not available.

Unbiased estimators of *R*^2^ have already been proposed in the literature for the in-sample estimator. In fact, the effect that adding predictors without predictive value has on the in-sample *R*^2^ is a well-known aspect in the regression literature, and the inflation (or bias) in the estimated value can be corrected with several techniques already available (see e.g. [Ezekiel, 1930a, Olkin and Pratt, 1958]). To the best of our knowledge, no correction is available in the literature for the out-of-sample case or when regularization is employed, while the effect on the out-of-sample measures has been recently studied for the mean square error [Rosset and Tibshirani, 2020].

Here we propose a correction for the bias terms introduced in the in-sample and out-of-sample *R*^2^ by overfitting and regularization. Our approach is based on approximations of quadratic forms. Our simulations show that the correction we propose is robust in different scenarios, and it allows recovering the correct 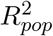. Importantly, our correction is identical to the adjusted *R*^2^ discussed in [Ezekiel, 1930b] for the in-sample estimate, and the results we obtain for the out-of-sample case bear resemblance to the case of the mean square error, which has been recently studied [Rosset and Tibshirani, 2020].

Our results are relevant for applications of linearized encoding in brain science in which models are compared on the basis of their explained variance. As it is well known in the field, comparing models based on an estimate of the explained variance in-sample is biased due to overfitting (i.e. in the case of two models with different dimensionality, even if the models have the same explanatory power, the model with the largest dimensions will have a larger *R*^2^). This is the reason why, following common practice in machine learning, applications have focused on comparing models using an estimate of the explained variance out-of-sample. Here we show that, in this case, the model with the largest dimensions will be more penalized by a larger *T*_*pess*_ component. That is, in the case of two models with same explanatory power, the model with the smallest dimensions will be erroneously chosen as the better model. It is important to note that our results indicate that, when comparing explained variance out-of-sample using an uncorrected estimator, if the largest model exhibits a higher predictive value, and it is thus selected as the better model, this result would not change when using an unbiased estimator as the one we propose here. However, previous model comparisons performed out-of-sample in which the smallest model resulted to have the largest explained variance should perhaps be re-examined using the correction we propose. Apart from the issue of selecting the better model, our results indicate that using a biased estimator (in or out-of-sample) results in an erroneous estimation of the *magnitude* of the difference in explained variance between models with different dimensions. Here we have also considered the case of regularization, and highlight the interplay between the error introduced by regularization and the one introduced by overfitting. Our results indicate that, when using regularization, the bias depends on the amount of regularization, an issue that further complicates previous model comparison attempts as regularization could be different for the different models. Importantly, our proposed correction is relatively insensitive to the amount of regularization, while it is important to stress that it is based on an approximation that may fail when extreme values of regularization are used. An interesting implication of our results is that, when using ridge regression, computational time could be saved by considering, for instance, a coarser grid of regularization values, since our proposed correction recovers the population *R*^2^ also for a sub-optimal regularization value.

Our simulations and the analyses of the real fMRI dataset suggest that the proposed 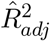 is an unbiased estimator of the true *R*^2^, which means that the expected value of 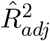 is equal to the population 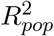. However, the adjustment can result, especially when used in-sample on regularized regression, in an inflation of the variance of the estimator, compared with the uncorrected in-sample or out-of-sample estimate, see simulations (see Figures 4 and 6). For model comparison purposes, an unbiased estimator with large variance should be preferred to a biased one with lower variance, as the latter would not control for false positives at the nominal level. However, large variance of an unbiased estimator could have consequences on statistical efficiency, resulting in a loss of power. This is less of a problem in the context of fMRI encoding studies, where the effects are generally evaluated on an area, and therefore the *R*^2^ estimates are averaged across a large number of voxels, reducing the variance by means of the averaging process.

Alternative strategies not considered in our work are based on performing model comparison by artificially equating the number of dimensions between models, either by reducing with Principal Component Analysis (PCA) the dimensions of the largest model or by adding noise dimensions to the smallest model. The idea behind the latter approach stems from the Ezekiel estimator, where the bias is linear with the number of dimensions. However, this strategy would be meaningful *only* in the in-sample OLS estimation, where the bias term is indeed a function of the dimensions *p* (see Table 1). In the out-of-sample, or even in the in-sample scenario with Ridge Regression, this is not anymore the case, and the covariance structure of the training (and testing) design matrix, together with the possible effect of regularization, make this approach inappropriate. Reducing the largest model with PCA is instead a better alternative, but introduces the issue of discarding variance explained by the largest model. If the reduction with PCA retains most of the variance, we expect this procedure to have similar results to our correction, while it would diverge if a larger portion of variance is discarded, hampering the predictive value of the (reduced) design matrix.

The statistical characterization of the *R*^2^ estimator has been at the core of recent investigations across several fields [Stijn Hawinkel and Maere, 2023] [Pospisil and Bair, 2021]. In particular, [Stijn Hawinkel and Maere, 2023] focuses on the out-of-sample *R*^2^, providing an unbiased estimator, together with confidence intervals obtained with a resampling procedure. It is important to note that [Stijn Hawinkel and Maere, 2023] focuses on a definition of *R*^2^ that is conditional to the number of samples, and the proposed approach is an estimate of the expected value across the distribution of datasets *of the same sample size*. This is consistent with the machine literature and it is desirable when validating a learning model versus chance. On the other hand, our approach is motivated by the need of decoupling the *R*^2^ estimate from the dimensionality of different feature spaces that may be compered on the basis of *R*^2^. That is, differently from [Stijn Hawinkel and Maere, 2023] we attempt to estimate 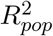. An interesting future direction may be to combine our framework with the procedure proposed in [Stijn Hawinkel and Maere, 2023] to estimate the confidence intervals of the *R*^2^ estimator we propose. In [Pospisil and Bair, 2021] the authors propose a correction that can be employed when comparing neural tuning curves with predictions based on models, conditional on the presence of multiple repetitions of the same experimental conditions. The authors show how an estimate of the variance across such repetitions can be used to account for biases in the *R*^2^ estimation. Although in their work there is not explicit emphasis on fitting models (the tuning curves are posited and compared with the neural recordings), some of the ideas presented could translate to the context of encoding models, provided that the same design is repeated several times. An interesting development in this direction may be to compare our estimator with the empirical one proposed in [Pospisil and Bair, 2021] in terms of design efficiency, studying which estimator has the lowest variance assuming a fixed duration of the experiment.

In a recent work, [Dupré la Tour et al., 2022] proposed an alternative framework for comparing models using a different metric, called “product metric”, in combination with Banded Ridge Regression. Although this procedure is more efficient than commonality analysis when the number of models to compare is larger than two, the metric used suffers from the same drawbacks as *R*^2^, as it can be shown that it falls back to the standard *R*^2^ estimate for in-sample OLS cases. In future work it may be relevant to investigate whether our framework extends to the product metric and can be used to account for the biasing effect of overfitting and regularization to produce an unbiased measure.

Throughout this work, we have made the assumption of independent and identically distributed (*i*.*i*.*d*.) noise term *ϵ*. Under this assumption, the use of OLS is optimal. However, when it is violated, different forms of regression should be employed (see Generalized Least Squares, [Diedrichsen and Shadmehr, 2005, Wicker and Fonlupt, 2003], which are not covered by our formulation. In the context of linearized encoding models, the validity of this assumption can be argued for the case of an encoding model based on beta estimates of single trials (or single stimuli), whereas it may need further validation when the regression is performed using the whole time series. Follow-up research may focus on determining whether, in the case of autocorrelated noise, the proposed corrections satisfactorily remove the bias terms in the *R*^2^ estimation, or whether modifications are needed.

## A Decomposition of *R*^2^ under Ridge Regression

By substituting the results of Eq (11) and Eq (12) in the definitions of 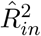 and 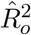 (Eq (2) and Eq (9)) we decompose of the explained variance in their contributing terms. For the in-sample *R*^2^, we have:

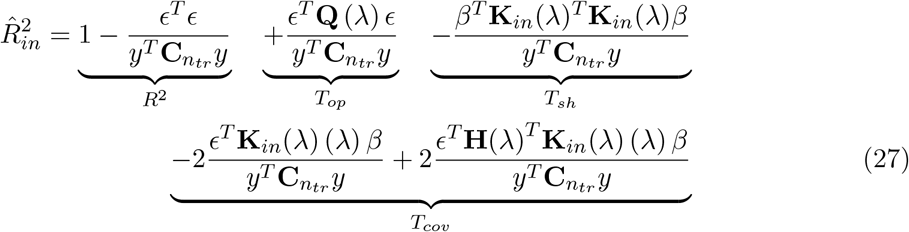

while for the out-of-sample 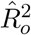 we have:

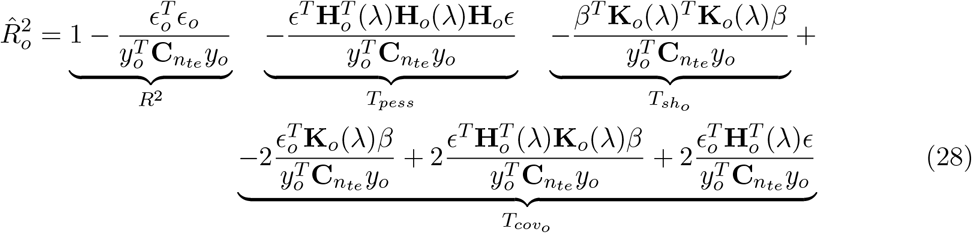

Importantly all terms involved in *T*_*cov*_ and 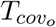 are cross-terms with zero expected value.

For this reason, their contribution to 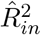 and 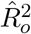 is minor compared to the contribution of the quadratic forms *T*_*op*_, *T*_*pess*_, *T*_*sh*_ and 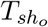.

## B Approximation of quadratic forms

Consider a symmetric positive semi-definite matrix **Q** and a vector **z. Q** is diagonalizable and we can write **Q** = **E**^*T*^ **DE** with **D** a diagonal matrix containing the eigenvalues *λ*_*i*_ and **E** is orthogonal. We can therefore express the quadratic form *u* = **z**^*T*^ **Qz**

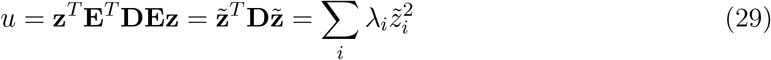

where 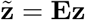 is a rotation of **z**.

We aim at linearizing the quadratic form *u* as a function of the L2 norm ||**z**||_2_, that is: *u* = *c*||**z**||_2_. Please note that, since **E** is an orthogonal matrix, it holds 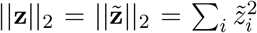.

We have then:

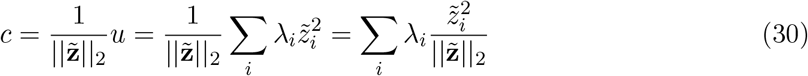

Note that *c* is a weighted average of the eigenvalues of **Q**. Considering the smallest and largest eigenvalues *λ*_*m*_ and *λ*_*M*_ of **Q**, we have:

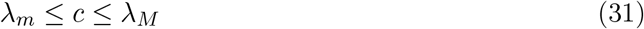

where the exact value of *c* depends on the components 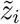 of 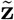, which is in general not available. We propose to approximate *c* as:

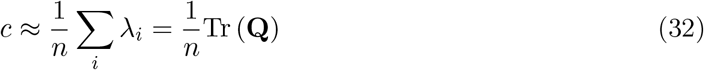

and therefore

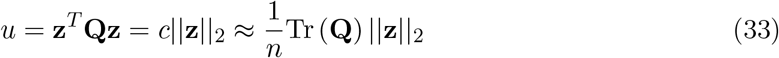

This approximation makes no explicit use of the unknown values of 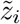. An intuition can be provided using a probabilistic argument: if we assume that 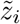 are random and *i*.*i*.*d*, then the expected value of all 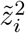 is equal and therefore the expected value of 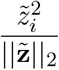 is 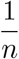.

## C Approximations for *T*_*op*_ and *T*_*pess*_

We describe here the approximations of *T*_*op*_ and *T*_*pess*_ used to correct the in-sample 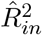 and the out-of-sample 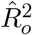 described in Section 2.6. Recalling Table 1, *T*_*op*_ and *T*_*pess*_ are defined as:

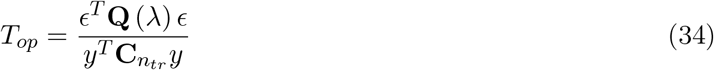

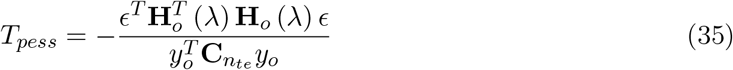

When computing *R*^2^ in-sample, we have from eq. (5) that 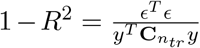. We can rewrite eq. (34) as:

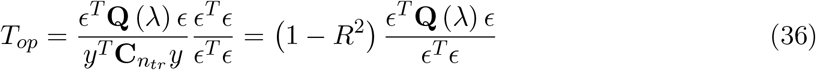

A similar argument can be used to show that, when estimating *R*^2^ out of sample, it holds:

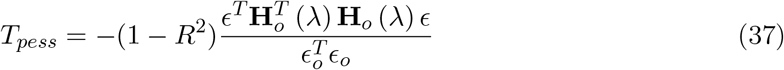

Please note that in the *T*_*op*_ term, we have the same noise component both at the numerator and denominator, while in the *T*_*pess*_ term, we have the noise of the training data on the numerator, and the noise of the testing data at the denominator. Furthermore, **Q** (*λ*) and **H**_*o*_ (*λ*) can be a function of *λ*, if regularization is used.

Using the approximation of quadratic forms presented in Appendix B, we can approximate *ϵ*^*T*^ **Q** (*λ*) *ϵ* as 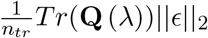 and *ϵ*^*T*^ *ϵ* as 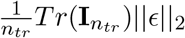, where 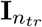 is the identity matrix of size *n*_*tr*_ *× n*_*tr*_. Noting that 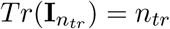, we have:

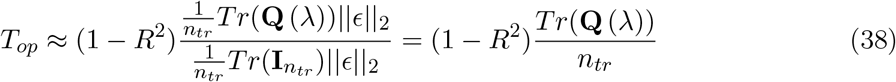

In the case OLS is used, this further reduces to 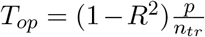, which leads to the Smith’s adjusted *R*^2^, as discussed in Section 2.6.

Applying the approximation of quadratic forms to the terms involved in *T*_*pess*_ (see eq. (37)), we have for the numerator: 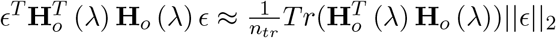 and for the denominator 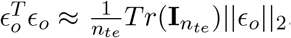. Therefore:

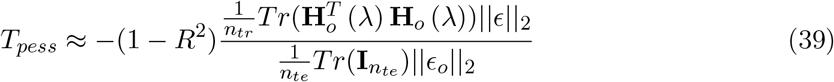

In this case, the two terms ||*ϵ*||_2_ and ||*ϵ*_*o*_||_2_ do not cancel each other out, as in eq. (38). However, we assume the same noise distribution for training and testing data, and therefore we can approximate the two terms with their expected value 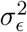. The pessimism term then becomes:

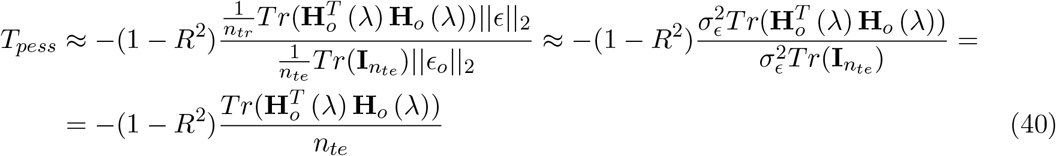

## D Approximation for *T*_*sh*_ and 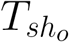

We will use the same approach presented in the previous section to approximate the bias terms due to shrinkage, *T*_*sh*_ for in the in-sample estimator and 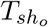 for the out-of-sample estimator. As shown in Section 2.5, the two terms can be written as a ratio of quadratic forms, see Eq. (16) and (17), which we rewite here for convenience:

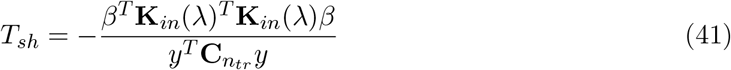

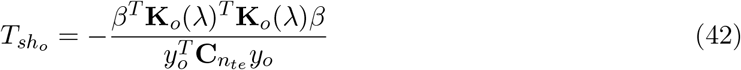

We start by approximating *R*^2^ as the ratio between the variance explained by the model and the total variance of the observation. Note that this approximation is exact only when using OLS in-sample, see Section 2.3. For the in-sample we have:

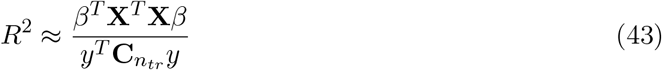

and for out-of-sample, we have:

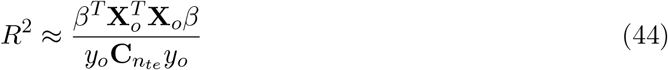

Combining eqs. (41) and (43), we have:

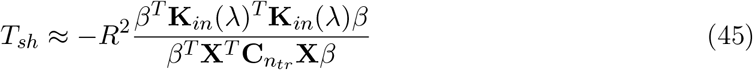

while for the out-of-sample, we have:

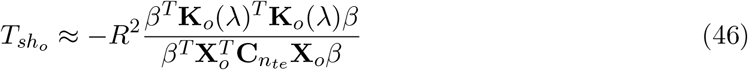

Approximating the quadratic forms at the numerator and denominator, as in Appendix C, we have:

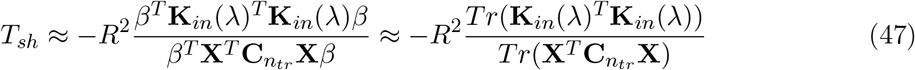

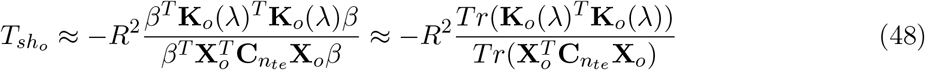

## References

Allen, E. J., St-Yves, G., Wu, Y., Breedlove, J. L., Prince, J. S., Dowdle, L. T., Nau, M., Caron, B., Pestilli, F., Charest, I., Hutchinson, J. B., Naselaris, T., and Kay, K. (2022). A massive 7t fmri dataset to bridge cognitive neuroscience and artificial intelligence. Nature Neuroscience, 25(1):116–126.

Bishop, C. M. (2007). Pattern Recognition and Machine Learning (Information Science and Statistics). Springer, 1 edition.

Chen, Q. and Qi, J. (2023). How much should we trust r 2 and adjusted r 2: evidence from regressions in top economics journals and monte carlo simulations. Journal of Applied Economics, 26(1):2207326.

Cichy, R. M., Khosla, A., Pantazis, D., Torralba, A., and Oliva, A. (2016). Comparison of deep neural networks to spatio-temporal cortical dynamics of human visual object recognition reveals hierarchical correspondence. Scientific Reports, 6.

Cohen, J. D., Daw, N., Engelhardt, B., Hasson, U., Li, K., Niv, Y., Norman, K. A., Pillow, J., Ramadge, P. J., Turk-Browne, N. B., et al. (2017). Computational approaches to fmri analysis. Nature neuroscience, 20(3):304–313.

De Angelis, V., De Martino, F., Moerel, M., Santoro, R., Hausfeld, L., and Formisano, E. (2018). Cortical processing of pitch: Model-based encoding and decoding of auditory fmri responses to real-life sounds. Neuroimage, 180:291–300.

de Heer, W. A., Huth, A. G., Griffiths, T. L., Gallant, J. L., and Theunissen, F. E. (2017). The hierarchical cortical organization of human speech processing. Journal of Neuroscience, 37(27):6539–6557.

De Martino, F., Moerel, M., van de Moortele, P.-F., Ugurbil, K., Goebel, R., Yacoub, E., and Formisano, E. (2013). Spatial organization of frequency preference and selectivity in the human inferior colliculus. Nature communications, 4(1):1386.

De Martino, F., Yacoub, E., Kemper, V., Moerel, M., Uludağ, K., De Weerd, P., Ugurbil, K., Goebel, R., and Formisano, E. (2018). The impact of ultra-high field mri on cognitive and computational neuroimaging. NeuroImage, 168:366–382. Neuroimaging with Ultra-high Field MRI: Present and Future.

Diedrichsen, J. and Shadmehr, R. (2005). Detecting and adjusting for artifacts in fmri time series data. NeuroImage, 27(3):624–634.

[Dupré la Tour et al., 2022]Dupré la Tour, T., Eickenberg, M., Nunez-Elizalde, A. O., and Gallant, J. L. (2022). Feature-space selection with banded ridge regression. NeuroImage, 264:119728.

Ezekiel, M. (1930a). Methods of Correlation Analysis. J. Wiley & Sons Incorporated.

Ezekiel, M. (1930b). The Sampling Variability of Linear and Curvilinear Regressions: A First Approximation to the Reliability of the Results Secured by the Graphic “Successive Approximation” Method. The Annals of Mathematical Statistics, 1(4):275–333. Publisher: Institute of Mathematical Statistics.

Gifford, A. T., Lahner, B., Saba-Sadiya, S., Vilas, M. G., Lascelles, A., Oliva, A., Kay, K., Roig, G., and Cichy, R. M. (2023). The algonauts project 2023 challenge: How the human brain makes sense of natural scenes.

Güçlü, U. and van Gerven, M. (2014). Deep neural networks reveal a gradient in the complexity of neural representations across the ventral stream. The Journal of Neuroscience, 35:10005 – 10014.

Hastie, T., Tibshirani, R., and Friedman, J. (2001). The Elements of Statistical Learning. Springer Series in Statistics. Springer New York Inc., New York, NY, USA.

Karch, J. (2020). Improving on adjusted r-squared. Collabra: Psychology, 6(1):45.

Kay, K. N., Naselaris, T., Prenger, R. J., and Gallant, J. L. (2008). Identifying natural images from human brain activity. Nature, 452:352–355.

Kell, A. J., Yamins, D. L., Shook, E. N., Norman-Haignere, S. V., and McDermott, J. H. (2018). A task-optimized neural network replicates human auditory behavior, predicts brain responses, and reveals a cortical processing hierarchy. Neuron, 98(3):630–644.

Kriegeskorte, N. and Douglas, P. K. (2019). Interpreting encoding and decoding models. Current Opinion in Neurobiology, 55:167–179. Machine Learning, Big Data, and Neuroscience.

Krizhevsky, A., Sutskever, I., and Hinton, G. E. (2012). Imagenet classification with deep convolutional neural networks. 25.

Kromrey, J. D. and Hines, C. V. (1995). Use of empirical estimates of shrinkage in multiple regression: a caution. Educational and Psychological Measurement, 55(6):901–925.

Lage-Castellanos, A., Valente, G., Formisano, E., and De Martino, F. (2019). Methods for computing the maximum performance of computational models of fmri responses. PLOS Computational Biology, 15(3):1–25.

LeBel, A., Wagner, L., Jain, S., Adhikari-Desai, A., Gupta, B., Morgenthal, A., Tang, J., Xu, L., and Huth, A. G. (2023). A natural language fmri dataset for voxelwise encoding models. Scientific Data, 10(1):555.

Marrazzo, G., De Martino, F., Lage-Castellanos, A., Vaessen, M. J., and de Gelder, B. (2023). Voxelwise encoding models of body stimuli reveal a representational gradient from low-level visual features to postural features in occipitotemporal cortex. NeuroImage, page 120240.

MathWorks (2020). 9.9.0.2037887 (R2020b). The MathWorks Inc., Natick, Massachusetts.

Moerel, M., De Martino, F., and Formisano, E. (2012). Processing of natural sounds in human auditory cortex: tonotopy, spectral tuning, and relation to voice sensitivity. Journal of Neuroscience, 32(41):14205–14216.

Moerel, M., De Martino, F., Santoro, R., Ugurbil, K., Goebel, R., Yacoub, E., and Formisano, E. (2013). Processing of natural sounds: characterization of multipeak spectral tuning in human auditory cortex. Journal of Neuroscience, 33(29):11888–11898.

Naselaris, T., Kay, K. N., Nishimoto, S., and Gallant, J. L. (2011). Encoding and decoding in fmri. NeuroImage, 56(2):400–410.

Naselaris, T., Olman, C. A., Stansbury, D. E., Ugurbil, K., and Gallant, J. L. (2015). A voxel-wise encoding model for early visual areas decodes mental images of remembered scenes. NeuroImage, 105:215–228.

Norman-Haignere, S. V. and McDermott, J. H. (2018). Neural responses to natural and model-matched stimuli reveal distinct computations in primary and nonprimary auditory cortex. PLoS biology, 16(12):e2005127.

Olkin, I. and Pratt, J. W. (1958). Unbiased Estimation of Certain Correlation Coefficients. The Annals of Mathematical Statistics, 29(1):201 – 211.

Pospisil, D. A. and Bair, W. (2021). The unbiased estimation of the fraction of variance explained by a model. PLOS Computational Biology, 17(8):1–36.

Raju, N. S., Bilgic, R., Edwards, J. E., and Fleer, P. F. (1997). Methodology Review: Estimation of Population Validity and Cross-Validity, and the Use of Equal Weights in Prediction. Applied Psychological Measurement, 21(4):291–305. Publisher: SAGE Publications Inc.

Rao, C. R. (1973). Linear statistical inference and its applications, volume 2. Wiley New York.

Rosset, S. and Tibshirani, R. J. (2020). From Fixed-X to Random-X regression: Bias-Variance decompositions, covariance penalties, and prediction error estimation. Journal of the American Statistical Association, 115(529):138–151.

Santoro, R., Moerel, M., De Martino, F., Goebel, R., Ugurbil, K., Yacoub, E., and Formisano, E. (2014). Encoding of natural sounds at multiple spectral and temporal resolutions in the human auditory cortex. PLoS computational biology, 10(1):e1003412.

Seibold, D. R. and McPhee, R. D. (1979). Commonality analysis: A method for decomposing explained variance in multiple regression analyses. Human Communication Research, 5(4):355–365.

Stijn Hawinkel, W. W. and Maere, S. (2023). Out-of-sample r2: Estimation and inference. The American Statistician, 0(0):1–11.

Storrs, K. R., Kietzmann, T. C., Walther, A., Mehrer, J., and Kriegeskorte, N. (2021). Diverse Deep Neural Networks All Predict Human Inferior Temporal Cortex Well, After Training and Fitting. Journal of Cognitive Neuroscience, 33(10):2044–2064.

Valente, G., Castellanos, A. L., Hausfeld, L., De Martino, F., and Formisano, E. (2021). Cross-validation and permutations in MVPA: Validity of permutation strategies and power of cross-validation schemes. NeuroImage, 238:118145.

Wicker, B. and Fonlupt, P. (2003). Generalized least-squares method applied to fmri time series with empirically determined correlation matrix. NeuroImage, 18(3):588–594.

Yin, P. and Fan, X. (2001). Estimating R 2 Shrinkage in Multiple Regression: A Comparison of Different Analytical Methods. The Journal of Experimental Education. Publisher: Taylor & Francis Group.

